# Intra-individual changes in the frequency of mosaic loss of chromosome Y over time estimated with a new method

**DOI:** 10.1101/631713

**Authors:** Marcus Danielsson, Jonatan Halvardson, Hanna Davies, Behrooz Torabi Moghadam, Jonas Mattisson, Edyta Rychlicka-Buniowska, Janusz Jaszczyński, Julia Heintz, Lars Lannfelt, Vilmantas Giedraitis, Martin Ingelsson, Jan P. Dumanski, Lars A. Forsberg

## Abstract

**Background:** Mosaic loss of chromosome Y (LOY) is the most common somatic mutation and is associated with all-cause mortality, non-haematological cancers and Alzheimer’s disease among other outcomes. The predominant method used for estimating LOY is the intensity data generated by SNP-arrays, which is difficult to interpret due to its logarithmic scale. Here we describe a new way to convert the LOY mosaicism into a non-logarithmic scale, which instead represents the percentage of affected cells.

**Methods:** We compared three independent LOY readouts from matched samples, generated by SNP-array, whole genome sequencing and droplet digital PCR. The SNP-array standardization was derived from this comparison and was applied in analyses of serially collected samples from a large cohort of aging men. The sampling was performed up to five times, spanning up to 22 years.

**Results:** We observed a higher correlation between the LOY measurements from SNP-array and the two other readouts when using the standardized, instead of the logarithmic, SNP-array data. We also observed a pronounced intra-individual variation of changes in the frequency of LOY within individual males over time.

**Conclusions:** Describing LOY measurements generated from SNP-arrays in percentage of cells without the Y chromosome makes comparisons to WGS and ddPCR measurements more precise and easier to interpret. This standardization could be applied to the vast amount of SNP-array data already generated in the scientific community, allowing further discoveries of LOY associated disease and outcomes. Additionally, the frequency of LOY in this study changed profoundly within men over time, likely as a result of aberrant clonal expansions.

## Background

Somatic mosaicism is defined as the presence of post-zygotic mutations in the soma of an organism. Mosaic loss of chromosome Y (LOY) refers to Y chromosome aneuploidy acquired during life and it is the most common post-zygotic mutation in human blood cells, affecting ∼1.6% of the genome(1). For over 50 years it has been known that LOY is a frequent event in cells of the hematopoietic system(2) and LOY in leukocytes was long viewed as a neutral event related to normal aging without phenotypical consequences(3). However, recent studies suggest the opposite as LOY has been found to be associated with increased risk for all-cause mortality(4, 5) as well as a growing list of diverse diseases and outcomes such as various forms of cancer(4, 6-9), autoimmune conditions(10, 11), Alzheimer’s disease(12), major cardiovascular events(13, 14), suicide completion(15), schizophrenia(16), diabetes(14) as well as age-related macular degeneration (AMD)(17).

At the single cell level LOY is a binary event, but it is manifested at the level of an individual as a gradual mosaicism, ranging from zero to 100% of cells without a Y chromosome. Recent studies have established that the frequency of LOY in leukocytes increases with age and that it occurs in about 5-10%, 15-20% and 20-30% of aging men around 60, 70 and 80 years of age, respectively(4, 7, 12, 18, 19). Furthermore, a recent study showed that the frequency of LOY in blood cells was 57% in 93 year old men(20). Although aging itself clearly is a very important risk factor, LOY in blood cells have also been reported in younger men(8, 17, 19) and other tissues (ectodermally-derived buccal mucosa), although in lower frequency than in haematopoietic cells(20). Thus, further studies are needed to fully characterize its prevalence, dynamic changes over time as well as potential phenotypic effects during the entire lifespan and across many tissues. Additional risk factors have been described and include smoking(7, 17, 18, 21), exposure to air pollution(22) as well as genetic background(7, 18, 23).

Measurements of LOY mosaicism from DNA have and could be performed using technologies such as karyotyping, qPCR, DNA-arrays and next generation sequencing (NGS) (Additional file 2: Supplementary Table 1). During the last decades, many millions of human DNA samples have been characterized in different large scale human genome projects and could readily be analyzed for occurrence of somatic structural variants and aneuploidies such as LOY in individual samples. For example, recent studies have reanalyzed data generated with various SNP-arrays, originally intended for genome wide association studies (GWAS) to estimate the level of LOY. The normalized intensity data captured by the array (Log R Ratio, LRR) reflect the DNA copy number in different regions of the genome. Hence, the measurement of LOY can be calculated from the median of the LRR values of the probes located within the male-specific region of chromosome Y (MSY, chrY: 2,781,480– 56,887,902, hg19/GRCh38.p12). This method typically generates an estimation for LOY called mLRRY (median Log R Ratio on male specific chromosome Y) where individuals without LOY display an mLRRY value around zero and a decreasing mLRRY value indicates an increasing level of LOY mosaicism. However, this inversed relationship is a shortcoming for intuitive interpretation of the mosaicism. To solve this problem, we here present a new method to transform the mLRRY data into a more intuitive unit, i.e. the percentage of cells with LOY (LOY%), which range between 0 and 100% and increases with the level of mosaicism. We also applied this transformation in comprehensive analyses of serially collected samples from aging men to characterize previously an unknown intra-individual variation of changes in the frequency of LOY within the blood of individuals studied over time.

## Results

We used data generated by SNP-array, whole genome sequencing (WGS) and droplet digital PCR (ddPCR targeting the *AMELX*/*AMELY* polymorphism) to estimate the level of LOY in DNA samples extracted from peripheral blood nucleated cells. The same DNA samples were analysed using these three methods and the estimated level of LOY in each sample from each technology is provided in Additional file 2: Supplementary Table 2. A detailed description of how LOY was estimated using each approach is provided in the Methods section. Briefly, for SNP-array data, a continuous variable was calculated from the Log R Ratio (mLRRY) as a median intensity value. For WGS, the frequency of cells with the Y chromosome present was estimated from the ratio between the read depth on chromosome Y in relation to the full genome. Finally, in the ddPCR we quantified the relative number of X and Y chromosomes by targeting a 6 bp polymorphism present between the *AMELX* and *AMELY* genes.

The generated data enabled us to compare the measurements of LOY among the three independent technologies (Fig. 1) and we found WGS and ddPCR to have the highest concordance in LOY estimation (Fig. 1 panels a and b). Specifically, in the samples analysed with these two independent technologies, a close to perfect correlation in LOY-estimation was achieved (R^2^=0.998, p< 2.2×10^-16^, Fig. 1 panels a and b). Comparing the level of LOY estimated in the samples analysed using SNP-array and WGS as well as in the samples analysed using SNP-array and ddPCR, also showed a high degree of concordance (Fig. 1 panels a, c and d). However, in contrast to the linear correlation between WGS and ddPCR readouts, comparing the level of LOY estimated using SNP-array with WGS or ddPCR showed non-linear relationships (Fig. 1 panels c and d). To increase comparability between SNP-array and other methods, we transformed the SNP-array data according to a new equation (LOY% = 100*(1-2^2mLRRY^)) as further described in the Methods section. The transformation was made by adjusting mLRRY data to the LOY estimates for the same samples using WGS and ddPCR and applying the method on the SNP-array data generated a linear relationship (Fig. 1 panels a, e and f).

**Figure 1.**
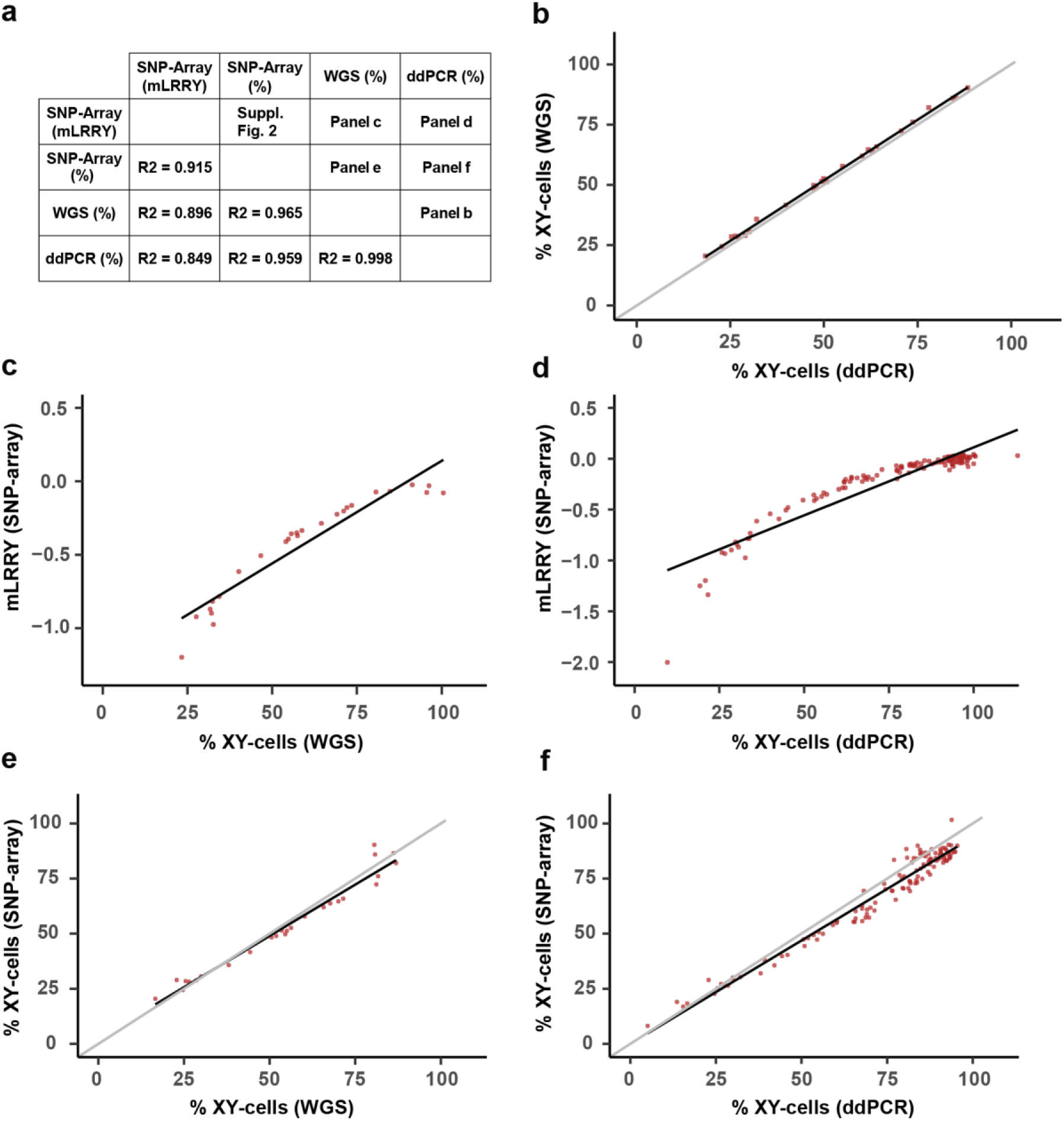
Comparisons between LOY measurements using three different methods, from the same set of DNA samples. The level of LOY was analysed using SNP-array, WGS and ddPCR in 121, 26 and 121 samples, respectively. By comparing the LOY measurements generated from the same sample using different technologies, we could estimate the accuracy for each method and the Pearson’s coefficient of determination for each correlation is presented in panel **a**. The Y-axes show the level of LOY estimated by different methods, represented as percentage of XY-cells (panels **b, e** and **f**) or as mLRRY (panels **c** and **d**). The X-axes in panels **b-f** show the percetange of XY-cells estimated with different technologies. In panels **b, e** and **f**, the grey lines represent the best theoretical fit and black lines show linear regressions. A linear correlation was observed between WGS and ddPCR (panels **a** and **b**). Non-linear relationships were observed between SNP-array and WGS (panel **a** and **c**) as well as between SNP-array and ddPCR (panels **a** and **d**). Transformation of SNP-array data increased the linearity of the relationship (panels **a, e** and **f**).

### Serial analyses of LOY in 276 aging men

The level of LOY from all available serially collected samples from the cohort Uppsala Longitudinal Study of Adult Men (ULSAM, www.pubcare.uu.se/ulsam) was estimated using Illumina SNP-array data, with both the conventional mLRRY as well as percent of cells with LOY as a metric. The serial analyses included data from 798 separate measurements of LOY in 276 men (median age = 81.9, range = 70-93) and each man was sampled 2-5 times over a period of up to 22.2 years. A main result from the serial analysis was an overall higher frequency of LOY within samples collected at higher ages (Fig. 2), confirming results of previous studies(4, 7, 12, 17-20). Furthermore, the serial analysis also revealed a previously undescribed profound inter-individual variation in the developmental trajectories of LOY clones in different men, i.e. variation in LOY driven aberrant clonal expansions (ACEs(1)), also referred to as clonal haematopoiesis (CH). For example, the result shows that in 67% of the studied individuals, the level of LOY did not change substantially during the study. We also found that the frequency of LOY increased during follow-up time in 26% of the individuals and decreased in 7% of the individuals (Fig. 2, Additional file 1: Supplementary Fig. 9 and Additional file 2: Supplementary Table. 3). More complex patterns could also be observed in a few individuals i.e. initial increase of LOY followed by a decrease but also initial decrease followed by an increase of LOY. These dynamic patterns were observed using both units to estimate LOY from the SNP-array data, i.e. mLRRY and percent of cells with LOY, respectively (Fig. 2 panels a and b).

**Figure 2.**
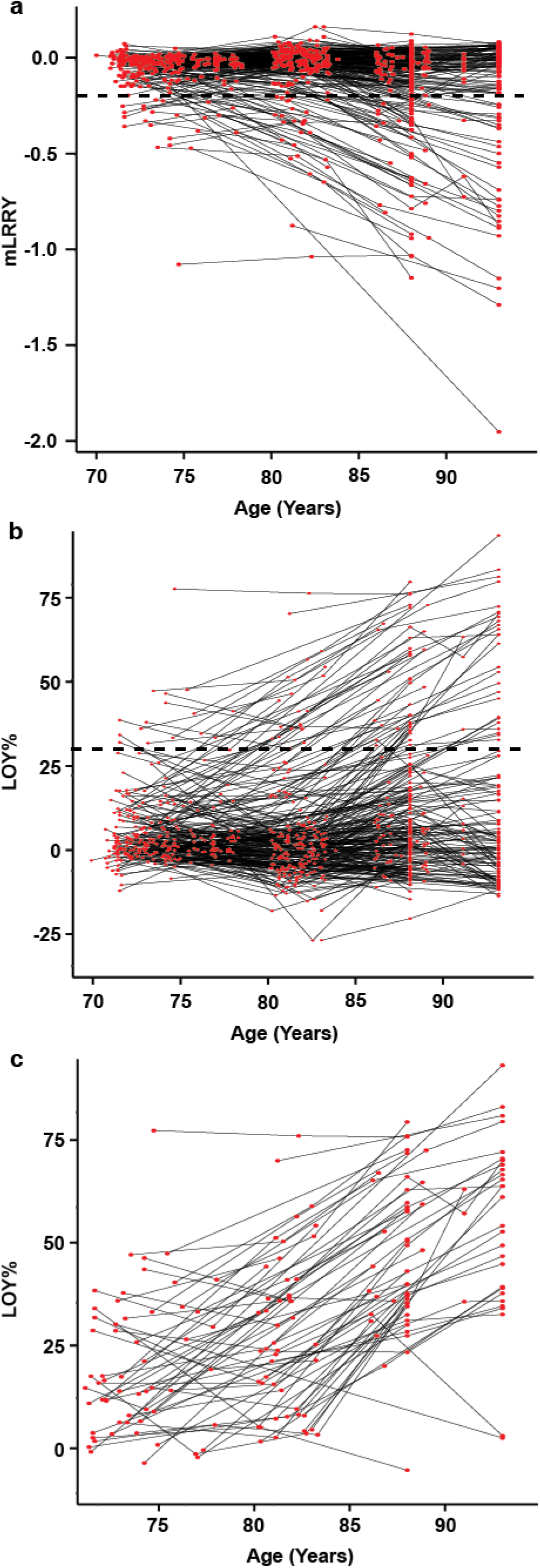
Results from serial analyses of LOY in whole blood DNA from 276 aging individuals sampled 2-5 times over a period of up to 22.2 years. X-axes show the age of sampling in years and Y-axes display the level of LOY estimated by SNP-array. In panel **a** the unit for LOY is the mLRRY and in panels **b** and **c** the unit is LOY%. Each red point represents a measurement of LOY in a single man and time point and black lines connects measurements from the same individual. Panels **a** and **b** visualise the changes in frequencies over time and dynamics of LOY clone evolution in blood within individually studied men. The dotted lines in panels **a** and **b** indicate a threshold at 30% LOY representing a high level of LOY and men with at least one measurement above the threshold are plotted in panel **c**.

As mentioned above, the frequency of LOY within a subset of the studied individuals showed a clear increase during follow-up time. The dotted lines in panels a and b in Fig. 2 mark a threshold where 30% of the studied cells in the samples are without the Y chromosome. Of the 276 men studied, we found that 65 individuals in at least one time point had a level of LOY on or above this threshold and their 183 measurements of LOY were plotted in Fig. 2 panel c. Within this group we observed that a subset of men showed a non-linear increase in frequency of LOY over time while others did not (Additional file 1:Supplementary Fig. 9). We tested for both linear and exponential associations between LOY and age in this group consisting of 183 data points and found that a linear predictor was stronger than the exponential (linear regression: R^2^=0.3442, p< 2.2×10^-16^ and exponential regression: R^2^=0.3298, p<2.2×10^-16^). Nevertheless, these group level analyses fail to reveal cases with a non-linear increase in frequency of LOY over time, which were observed in certain individuals (Fig. 2 and Additional file 1: Supplementary Fig. 9).

## Discussion

Measurements of LOY from DNA have been performed using many different technologies such as various types of karyotyping, qPCR, genotyping arrays and next generation sequencing (Additional file 2: Supplementary Table 1). Recent studies have reanalyzed data generated with various SNP-arrays, originally intended for genome wide association studies (GWAS), to estimate the level of LOY in individuals and described profound phenotypic effects associated with LOY in leukocytes. We here evaluated the three independent technologies SNP-arrays, WGS and ddPCR (targeting the *AMELX*/*AMELY* polymorphism) for measuring LOY mosaicism. We analysed the same DNA samples using these three methods which enabled comparisons of the level of LOY estimated by the different technologies.

We established that LOY estimation using WGS and ddPCR yielded close to identical results from the same DNA samples tested. However, the corresponding comparisons between SNP-array and WGS or ddPCR showed non-linear relationships, likely as an effect of the logarithmic scale of the intensity data generated by the SNP-array platforms. After scaling the SNP-array data using a new method, the comparisons to other methods increased linearity (Fig. 1). In addition, the unit percentage of cells with LOY, generated by the transformation, represents the studied biological event in a more straightforward way, since a higher percentage represents a higher level of mosaicism.

In order to evaluate the new method and the LOY% unit, we studied serial changes in the level of LOY mosaicism in DNA samples collected serially in a unique cohort of aging men called ULSAM. To our knowledge, this is the first study showing comprehensive LOY analyses from samples collected serially from the same men. The participants of the ULSAM study have been followed clinically for 48 years and blood samples have been collected repeatedly from the same participants. The serial sampling allowed us to, for the first time, study changes in the level of LOY within individuals over time and we found three main patterns. First, in a large part of the studied men, the levels of LOY were relatively low during the entire study period. Second, in other men the frequency of LOY increased substantially during follow-up time and third, in a few men we found that the level of LOY showed an initial increase followed by a decrease in frequency (Fig. 2 and Additional file 1: Supplementary Fig. 9). Thus, we validated results from previous studies showing that LOY is more frequent in older men(4, 7, 12, 18-20) and in addition, we described a novel finding of profound inter-individual variation in development of LOY clones in blood (Fig. 2). Interestingly, we observed in a subset of the studied men that the frequency of LOY increased in a non-linear rate (Additional file 1: Supplementary Fig. 9). These results illustrate the dynamic nature of aberrant clonal expansions (ACEs(1)) with LOY in the hematopoietic system. The non-linear increase in frequency of LOY over time is likely an effect of oligo or polyclonal processes, i.e. several hematopoietic progenitor cells giving rise to ACEs with LOY.

Here we describe a new method for transformation of LOY data generated by SNP-array and show its advantages applied on a data set where LOY was estimated on serially collected blood samples. The approach to re-analyze SNP-array data could be applied to the millions of experiments already generated in GWAS studies, to further investigate associations between LOY and various diseases and other phenotypic outcomes.

## Conclusions

Here we describe a new method for standardization of LOY data generated by SNP-array and show its advantages when comparing with LOY estimates using WGS or ddPCR. Furthermore, when applied on a data set where LOY was estimated on serially collected blood samples, data was easier to interpret with the intuitive scale and unit (LOY%). This standardization could be applied to estimate LOY in the millions of SNP-array experiments already generated in GWAS studies to further investigate associations between LOY and various diseases and other phenotypic outcomes.

## Methods

### Samples and DNA extraction

DNA was extracted from blood samples of participants in the Uppsala Longitudinal Study of Adult Men (ULSAM, www.pubcare.uu.se/ulsam) using the QIAamp DNA Blood kit (51194, Qiagen) according to the manufacturer’s instructions.

### SNP-array

The mLRRY value was calculated as the median of the Log R Ratio (LRR) value of each probe in the male specific Y (MSY) region of chromosome Y between the pseudoautosomal region 1 and pseudoautosomal region 2 (PAR-1 and PAR-2). The four different Illumina SNP-array platforms used in this study and the number of probes in the MSY and PAR regions for the genotyping platforms used in this study are described in Additional file 2: Supplementary Table 4. The mLRRY value calculated for each sample (N=121) was adjusted for batch effects using the positive tail of the distribution of mLRRY values for each batch as an estimator of the batch specific noise as previously described(4).

### mLRRY transformation

Transformation from mLRRY generated from MSY to percentage of cells was done in three steps. First mLRRY was antiloged (2^mLRRY^) and correlated to data from the same samples generated by WGS (N=26) or ddPCR (N=121). Secondly, a power equation was calculated from both correlations and independently of each other resulted in the same formula: 0.9*mLRRY^1.8^ (R2=0.97). Finally the formula was rounded to the nearest integer and used to adjust the antilog mLRRY, resulting in the following equations:

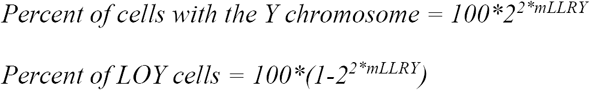

In parallel, data generated from the PAR-region from chromosome Y and X (median B-allele frequency, BAF) is presented in Additional file 1: Supplementary Figure 4 panel b, and was calculated accordingly(24):

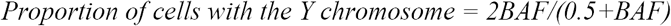

The data describing the proportion of normal cells calculated from B-allele frequency was also correlated to WGS and ddPCR and by rounding the resulting power equation to the nearest integer the relationship to percentage of cells could be described:

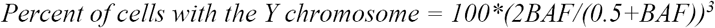

### Whole genome sequencing (WGS)

Sequencing libraries were prepared using the truseq Nano DNA sample preparation kit (T FC-121-4001/4002, Illumina Inc) extracting 100 ng DNA for each sample. Sequencing libraries were run on an Illumina HiSeq X instrument (version 2.5 sequencing chemistry) and sequenced to a depth of 30x. Each sequenced library had a read length of 150 bp with an insert size of 350 bp.

Sequencing reads were aligned to the GRCh37 human reference genome with the BWA aligner (version 0.7.12). Copy number for chromosome Y was estimated by the Control-Freec software using read counts in non-overlapping windows across the genome. These were fitted by the GC content and mappability information and the median ploidy for the Y chromosome was calculated(25).

### ddPCR

Bio-Rad’s QX200 Droplet Digital PCR System was used for the processing and fluorescent measurements of droplets. Additionally, Bio-Rad’s software QuantaSoft (version 1.7.4.0917) was used in both data generation and analysis. Extracted DNA samples with concentrations ranging between 300ng/µl to 20ng/µl were pre-digested for 15 minutes in 37°C with HindIII (Thermo Fischer, article number: #FD0504) and diluted with an equal volume of water. Subsequently 50 ng of the digested and diluted DNA sample was mixed in PCR supermix for probes without dUTP (BioRad, article number: 186-3023) together with primers and probes (Thermo Fisher, article number: C_990000001_10). PCR conditions used was an initial denaturation at 95°C for 10 minutes followed by 40 cycles of 94°C denaturation for 30 seconds and combined annealing and extension at 60°C for 1 minute. The PCR program ends with 98°C for 10 minutes and finally a 10°C hold. The fluorophores for the TaqMan probes are FAM for *AMELY* and VIC for *AMELX* and the schematics of the *AMELX*/*AMELY* TaqMan-assay used in this study is described in Additional file 1: Supplementary Figure 5. Samples were run in duplicates and if the standard deviation for the *AMELY*/*AMELX* ratio exceeded 1.2, it was re-run.

To estimate the limit of detection (LOD) for LOY, a dilution series was generated by mixing male and female DNA and determined by linear regression accordingly:

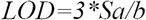

*Where Sa is the standard deviation and b is the slope of the dilution series.*

Pairwise two-tailed student t-test was used for comparing the control male DNA to each of the different steps in the dilution series. The measured *AMELY*/*AMELX* ratio was adjusted for any LOY in the male control DNA. Additionally, the difference between male and female genomic weight was also adjusted for when producing the dilution series.

## Declarations

### Ethics approval and consent to participate

The study had been approved by the Regional Ethical Committee in Uppsala, Sweden. All study participants provided written informed consent for participation. The reference number of the approvals are: Dnr: 02-018 approved 2002-06-05 (ethics for the early ULSAM genetics), dnr: 02-605 approved 2003-02-18 (ULSAM82), dnr: 2007/338 approved 2008-01-23 (ULSAM88) and dnr: 2013/350 approved 2013-10-23 (ULSAM93).

### Availability of data and material

The data generated in this study is either available (Additional file 2: Supplementary Table 2) or could be made available by request to corresponding authors.

### Authors’ contributions

MD, ERB, LL, VG, JJ, MI, JPD and LAF conceived the study. MD, HD, JPD and LAF designed the study. MD, HD, JH1 & JH2 performed the experiments. MD, JH1, BTM and JM analysed and interpreted the data. MD and LAF wrote the manuscript with input from all other authors. All authors have read and approved the final manuscript.

## Acknowledgements

We acknowledge K. Ström for collection of samples from the ULSAM. Thanks goes out also to U. Landegren for discussions and feedback on the manuscript.

## Supplemental material

**Supplementary figure 1.**
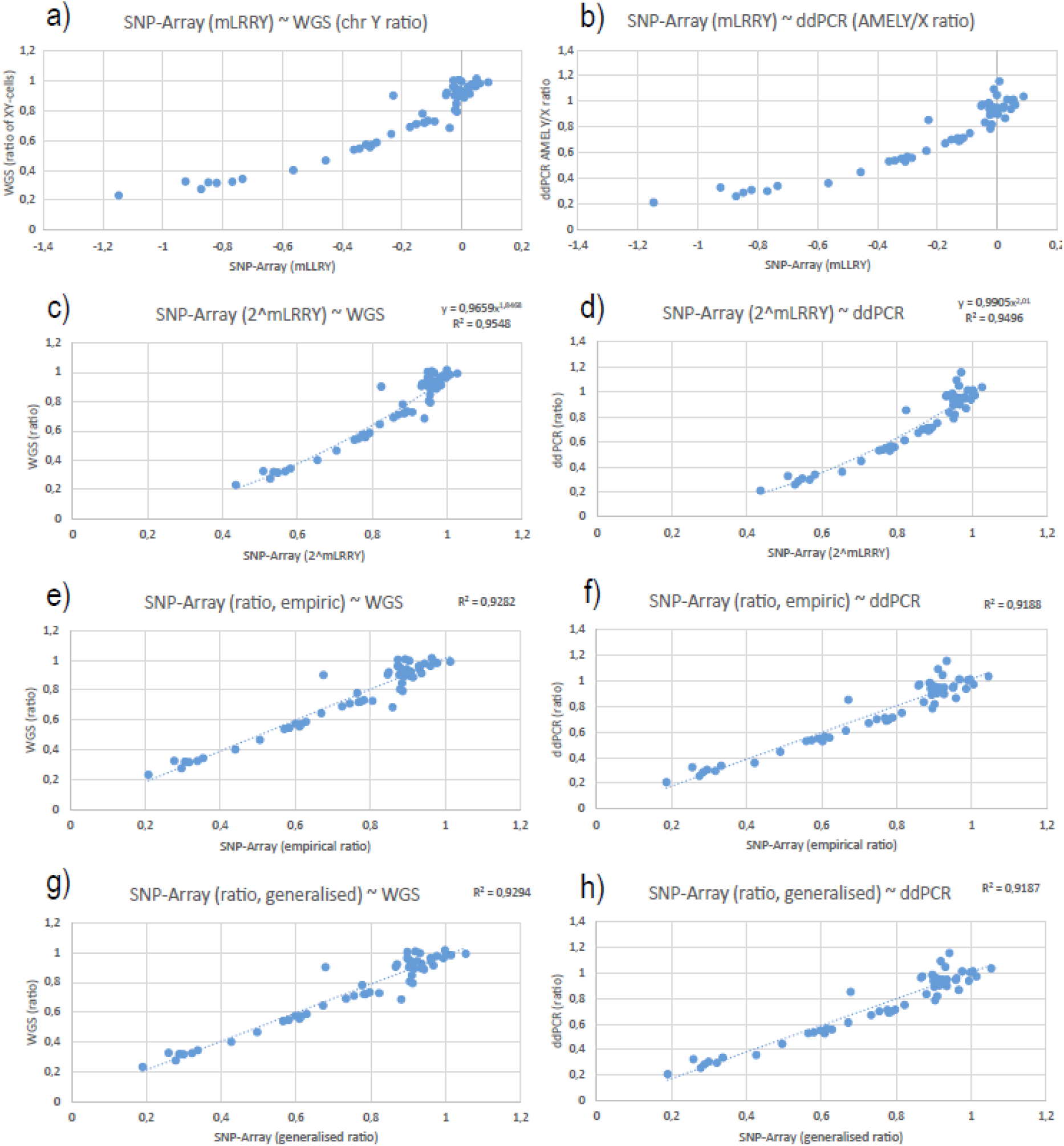
Measurement of LOY using three different methods. Measurements for mosaic loss of chromosome Y (LOY) using SNP-array, whole genome sequencing (WGS) and droplet digital PCR (ddPCR) with the *AMELY*/*AMELX*-assay from paired individuals (N=26). The plotting of SNP-array data (mLRRY) on the x-axis and either WGS median ratio of chromosome Y reads on the y-axis **(a)** or *AMELY*/*AMELX*-ratio from ddPCR on the y-axis **(b)** results in a problematic comparison due to non-linearity and difference in the scale. Antilog of SNP-Array mLRRY-data on the x-axis makes it possible to apply a power trendline with either *AMELY*/*AMELX*-ratio from ddPCR **(c)** or WGS median ratio of chromosome Y reads on the y-axis **(d)**. The SNP-array antilog mLRRY-data was adjusted according to the formula from the power trendline into an empirically determined ratio and plotted on the x-axis with the corresponding WGS ratio **(e)** or ddPCR data **(f)** on the y-axis. The formula for adjusting SNP-Array mLRRY to ratio of XY-cells was finally rounded to the closest integer and applied on the mLRRY data on the y-axis while comparing with LOY estimates generated by WGS **(g)** or ddPCR **(h)** plotted on the y-axis.

**Supplementary figure 2.**
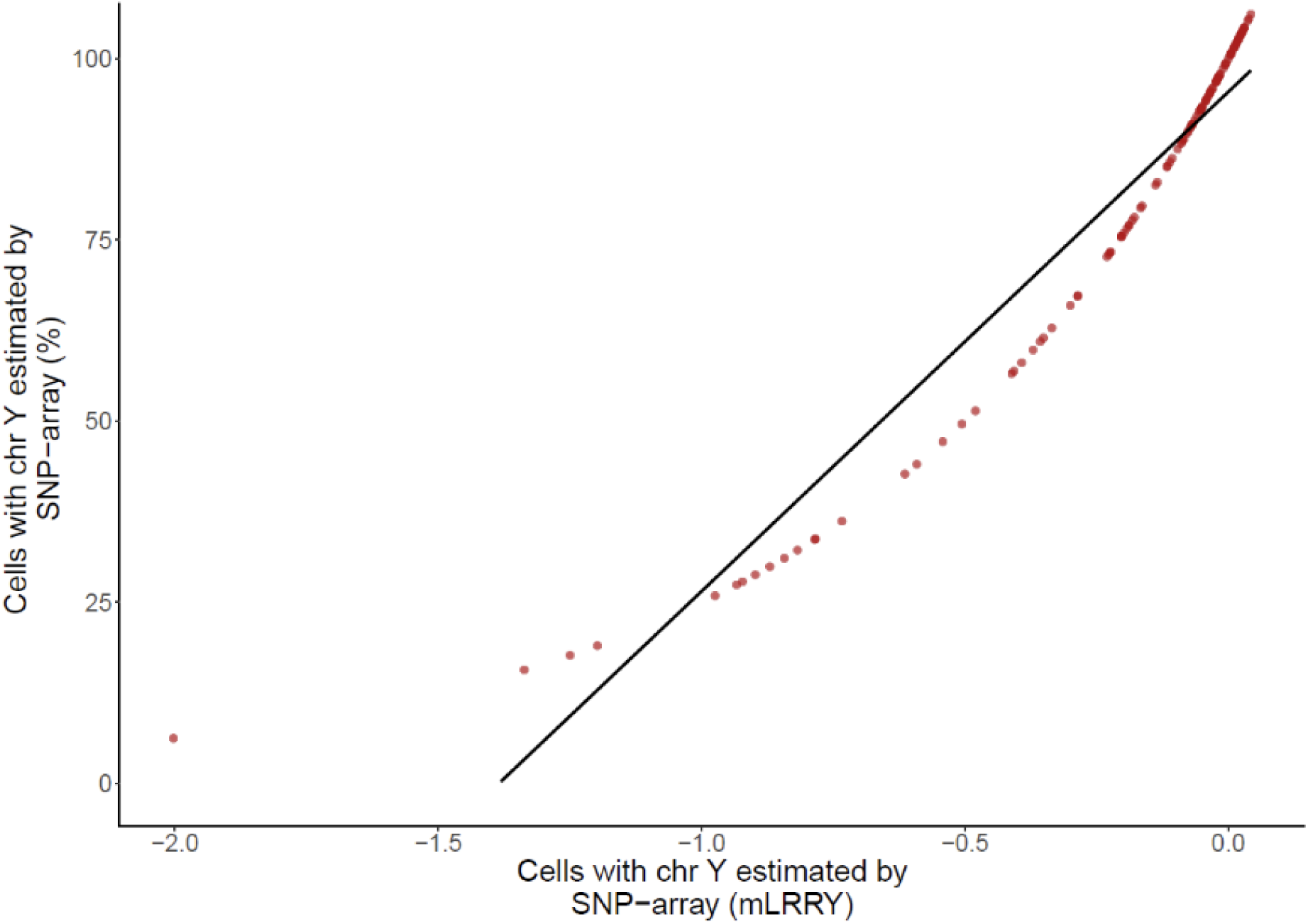
Correlation between two units of LOY measurements using SNP-Array (N=121) SNP-array data visualised on the x-axis by median Log R Ratio of the male specific chromosome Y (mLRRY) and on the y-axis as percent of normal cells generated using the transformation described in Methods.

**Supplementary figure 3.**
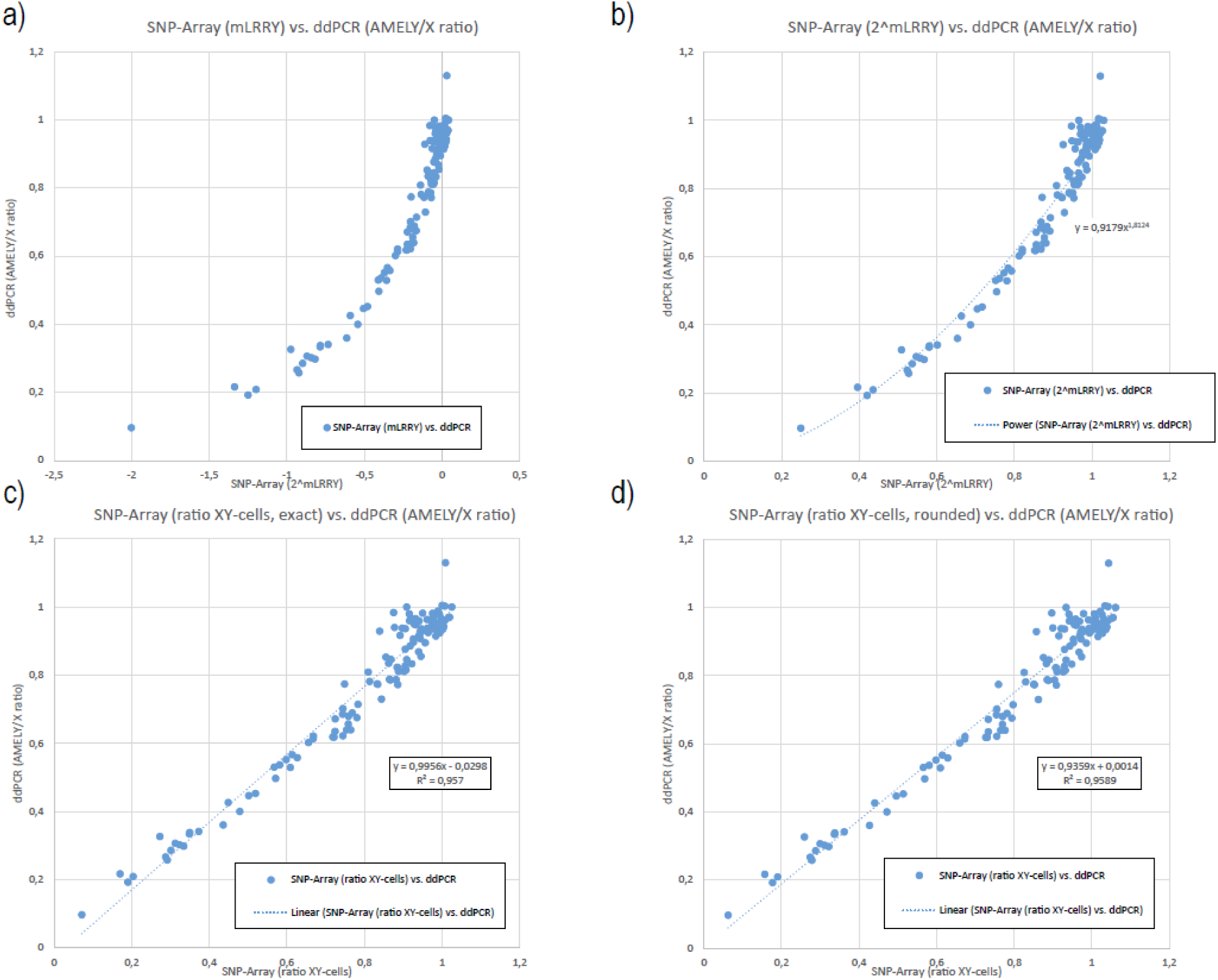
Correlation between *AMELY*/*AMELX* ratio generated from ddPCR to stepwise transformed SNP-Array data generated from the male specific chromosome Y (N=121) Ratio between *AMELY* and *AMELX* generated from ddPCR is plotted on the y-axis in all four plots and compared with data generated from SNP-Array from paired individuals (N=121) on the x-axis. Comparison using mLRRY **(a)** is again problematic with a non-linear correlation and both negative and positive values. Antilog of mLRRY (2^mLRRY) results in only positive values and makes it possible to apply a power trendline **(b)**. The generated power trendline formula was used to adjust the mLRRY **(c)**, resulting in a linear correlation between ddPCR and SNP-Array data. Adjusting mLRRY with a formula rounded to the closest integer in the power trendline formula **(d)** also resulted in a linear correlation but with an even higher correlation coefficient than the linear model using the exact values of the power trendline in panel **b**.

**Supplementary figure 4.**
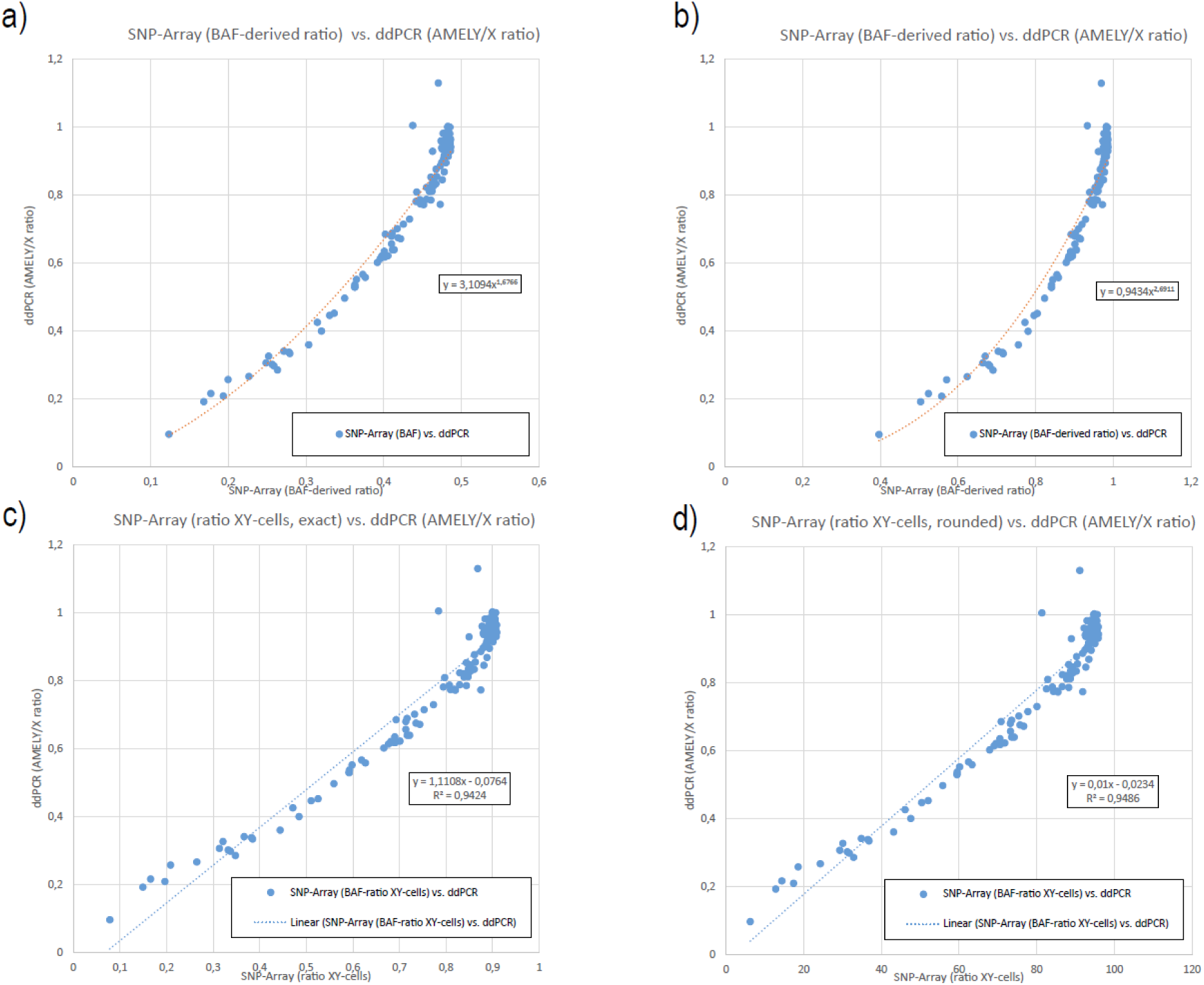
Correlation between AMELY/X ratio generated from ddPCR to stepwise transformed SNP-Array data generated from the pseudoautosomal region 1 of chromosome X and Y (N=121) The SNP-Arrays also generates data for the b-allele frequency (BAF) of the pseudoautosomal region 1 (PAR1). This can be used to study LOY as PAR1 is present on both the X and the Y chromosome. Ratio between AMELY and AMELX generated from ddPCR is plotted on the y-axis in all four plots and compared with data generated from SNP-Array from paired individuals (N=121) on the x-axis. Comparison using BAF **(a)** is again problematic due to non-linear correlation and spans only the region between 0 and 0.5. Using a previously published algorithm, the spanned region is increased to reach close to 1, but the non-linearity is unaffected **(b)**. The generated formula from the power trendline in **(b)** was used to adjust the BAF-derived data **(c)**, resulting in a linear correlation between ddPCR and SNP-Array data. Adjusting the BAF-derived data with the same formula but rounded to the closes integer **(d)** also resulted in a linear correlation but with an even higher correlation coefficient.

**Supplementary figure 5.**
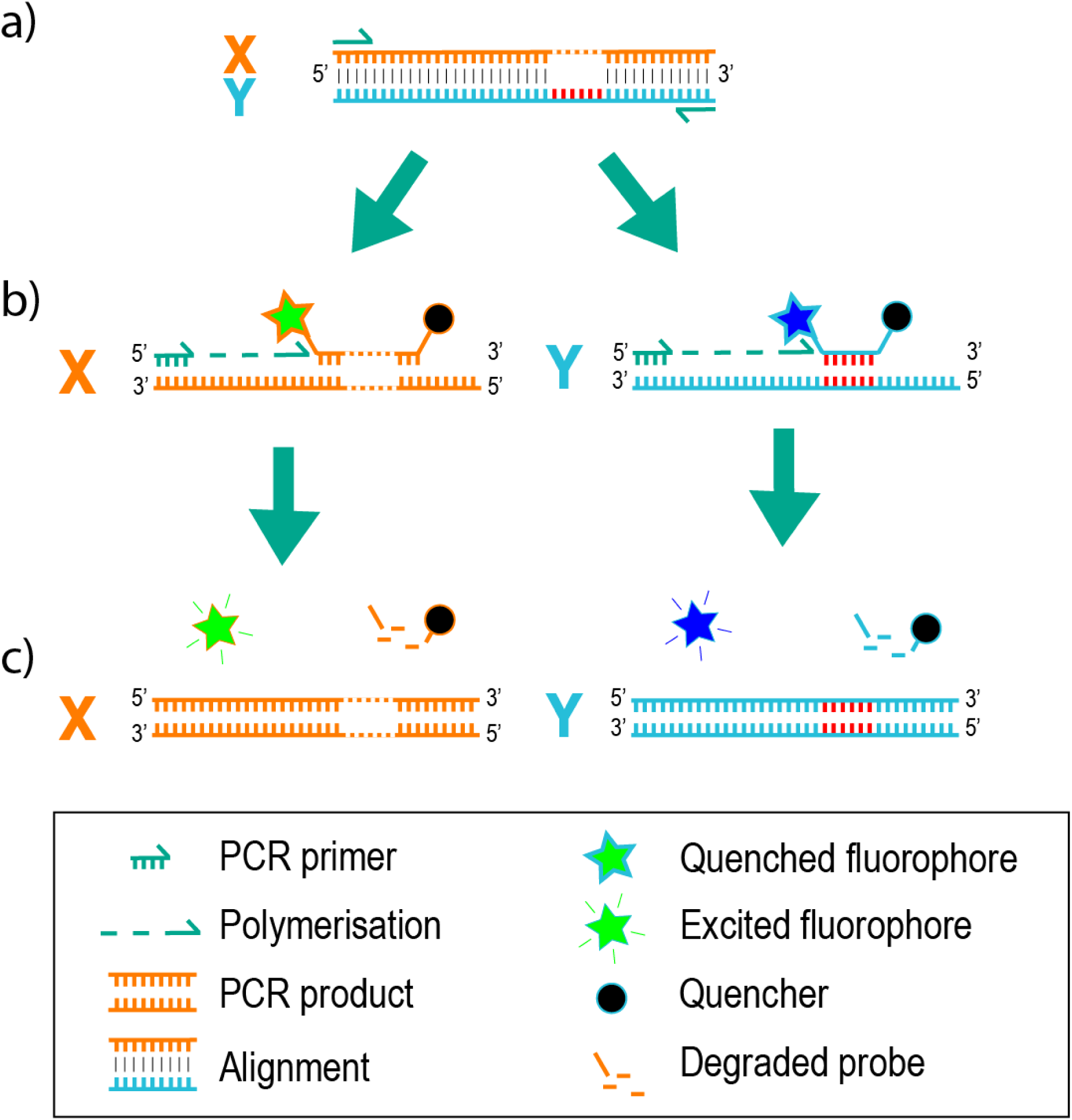
Schematics of the ddPCR assay *AMELX*/*AMELY*. Schematics of the *AMELX*/*AMELY* ddPCR taqman assay. The genetic target for this assay is the homologous genes *AMELX* and *AMELY* on chromosome X and Y respectively, that carries a 6-bp deletion in AMELX. **a)** Alignment of the region containing this 6-bp deletion that is targeted for PCR-amplification using identical forward and reverse primers for *AMELX* and *AMELY*. **b)** During the PCR-amplification, fluorescently labelled (and quenched) TaqMan probes specifically hybridise to either the deleted site on *AMELX* or the inserted site on AMELY. **c)** The fluorophores are cleaved of by the polymerase, allowing excitation of the fluorophore (FAM for *AMELY* and VIC for *AMELX*).

**Supplementary figure 6.**
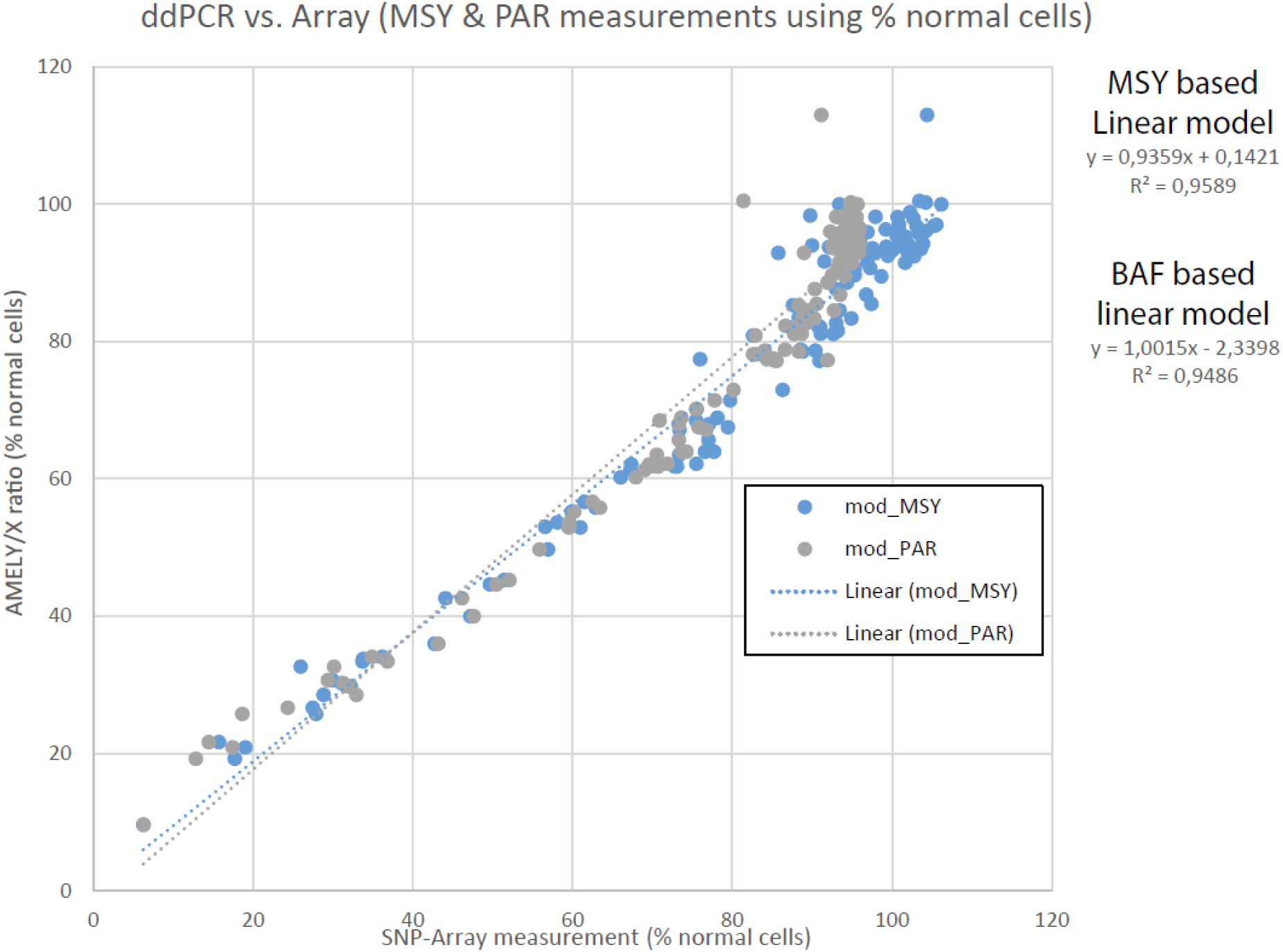
Correlation plot with ddPCR (percent *AMELY*/*AMELX*) and transformed SNP-Array data (MSY and BAF derived into percent XY-cells) Ratio between *AMELY* and *AMELX* expressed as percent normal-cells generated from ddPCR is plotted on the y-axis and compared with data generated from SNP-Array from paired individuals (N=121) on the x-axis. Data generated from MSY (transformed mLRRY) is plotted in blue and data generated from PAR1 (transformed BAF-derived data described in detail in Methods) is plotted in grey. The PAR-derived data highly correlates to the MSY-data except for the region over 90% XY-cells, which is reflected in a lower Pearson’s R^2^ of the linear model for PAR-derived compared to MSY-based data.

**Supplementary figure 7.**
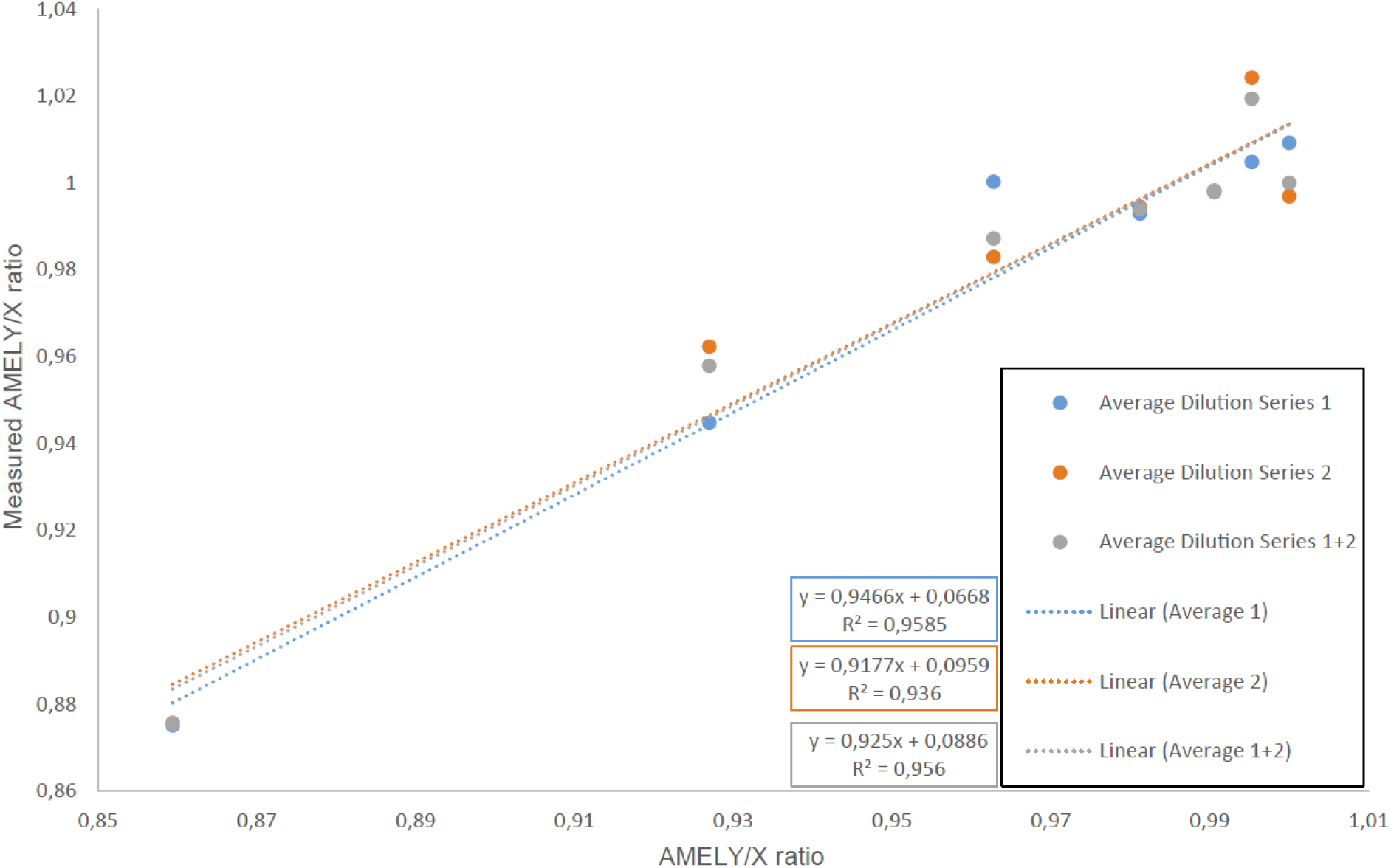
Linear model of dilution series measured with the *AMELY*/*AMELX* ddPCR-assay. The ddPCR-generated AMELY/X-ratio is plotted on the y-axis and the expected AMELY/X-ratio from the dilution of male and female control DNA is plotted on the x-axis. Blue dots represent the average from 4 replicates of each dilution step (Average dilution series 1). Orange dots represent the average from 12 separate replicates from the same dilution series (Average dilution series 2). The average from both dilution series is plotted in grey (Average dilution series 1+2). Linear regressions follow the same colour coding as the three series.

**Supplementary figure 8.**
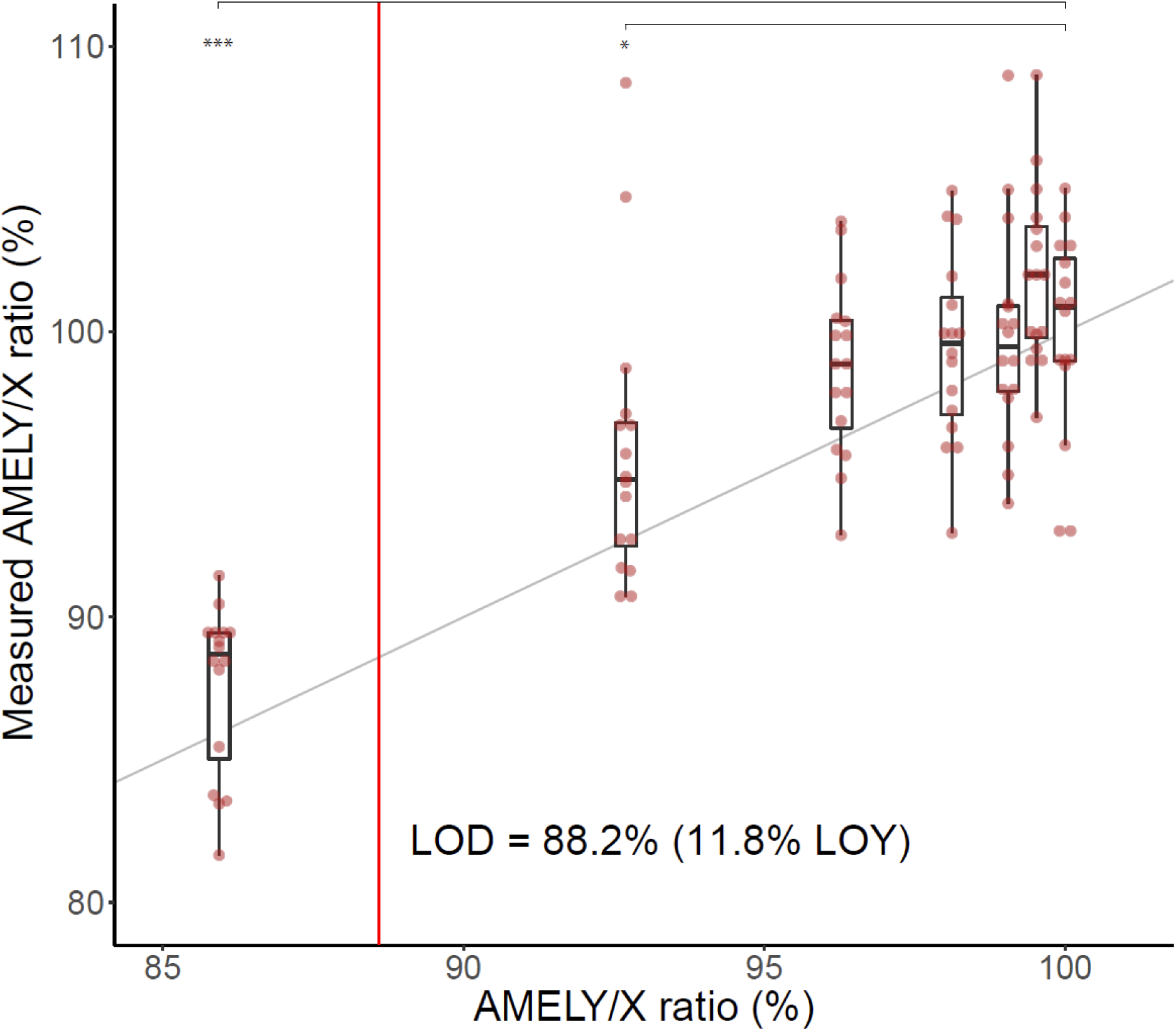
Analysis of limit of detection of LOY using the *AMELY*/*AMELX* polymorphism and ddPCR. Dilution series of chromosome Y, measured with the *AMELY*/*AMELX* ddPCR-assay. On X-axis are six different known levels of LOY made from mixing male & female DNA and on Y-axis are the measured *AMELY*/*AMELX* ratio. Limit of detection (LOD) was determined by linear regression by the formula 3xSD/b, also explained in methods. The data required for this calculation is disclosed in Supplementary Table 5. The lower hindge corresponds to the 25^th^ percentile and the upper hindge corresponds to the 75^th^ percentile. The minimum value of the lower whisker is 1.5 x IQR (inter quartile range) of the lower hindge and the maximum of the upper whisker is 1.5 x IQR of the upper hindge. Additionally, significance level * indicates P<0.05 and *** indicates P<0.0001, from a pairwise student t-test (two-sided) where every dilution step was compared to the same male control DNA (*AMELY*/*X* ratio = 100%).

**Supplementary figure 9.**
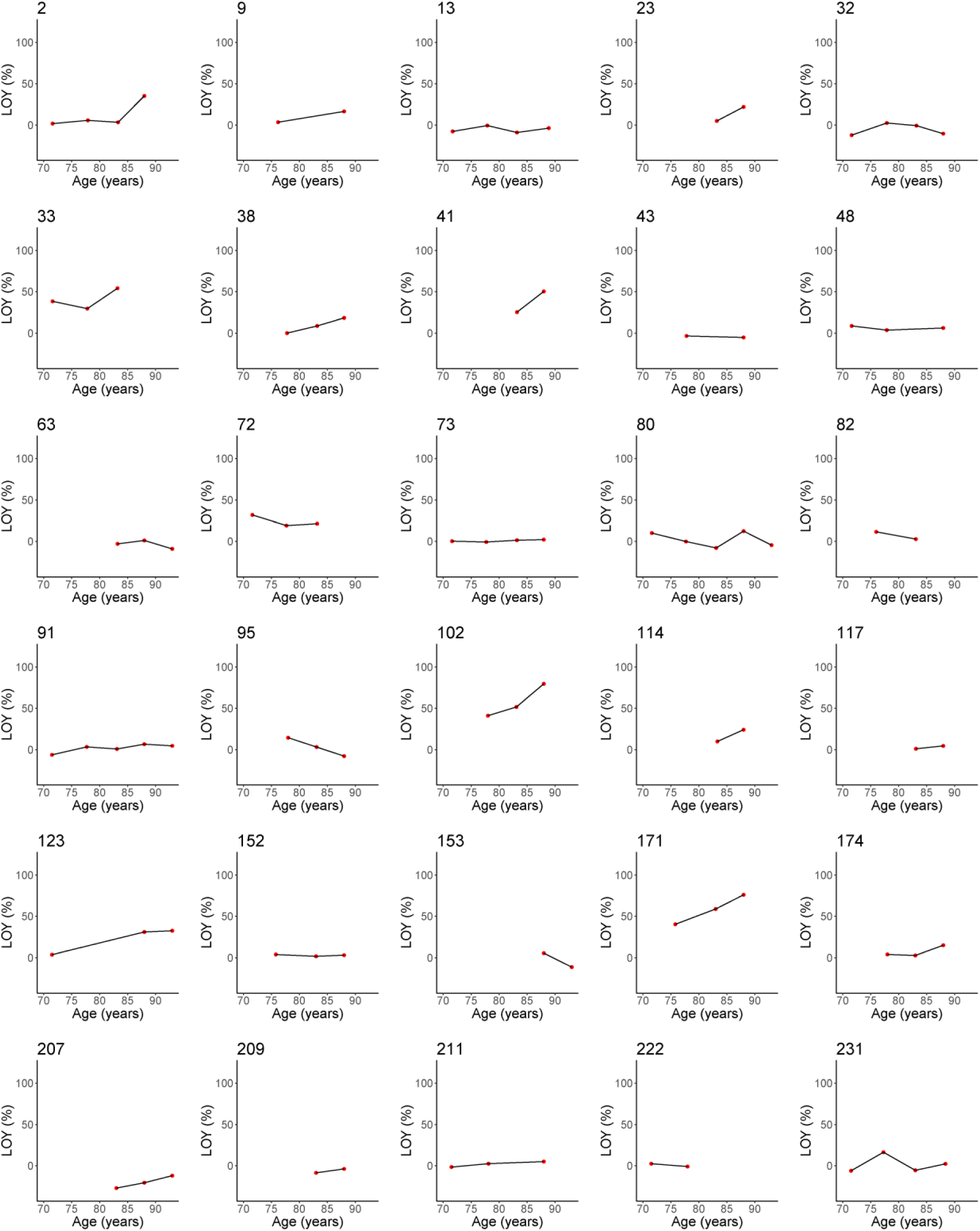
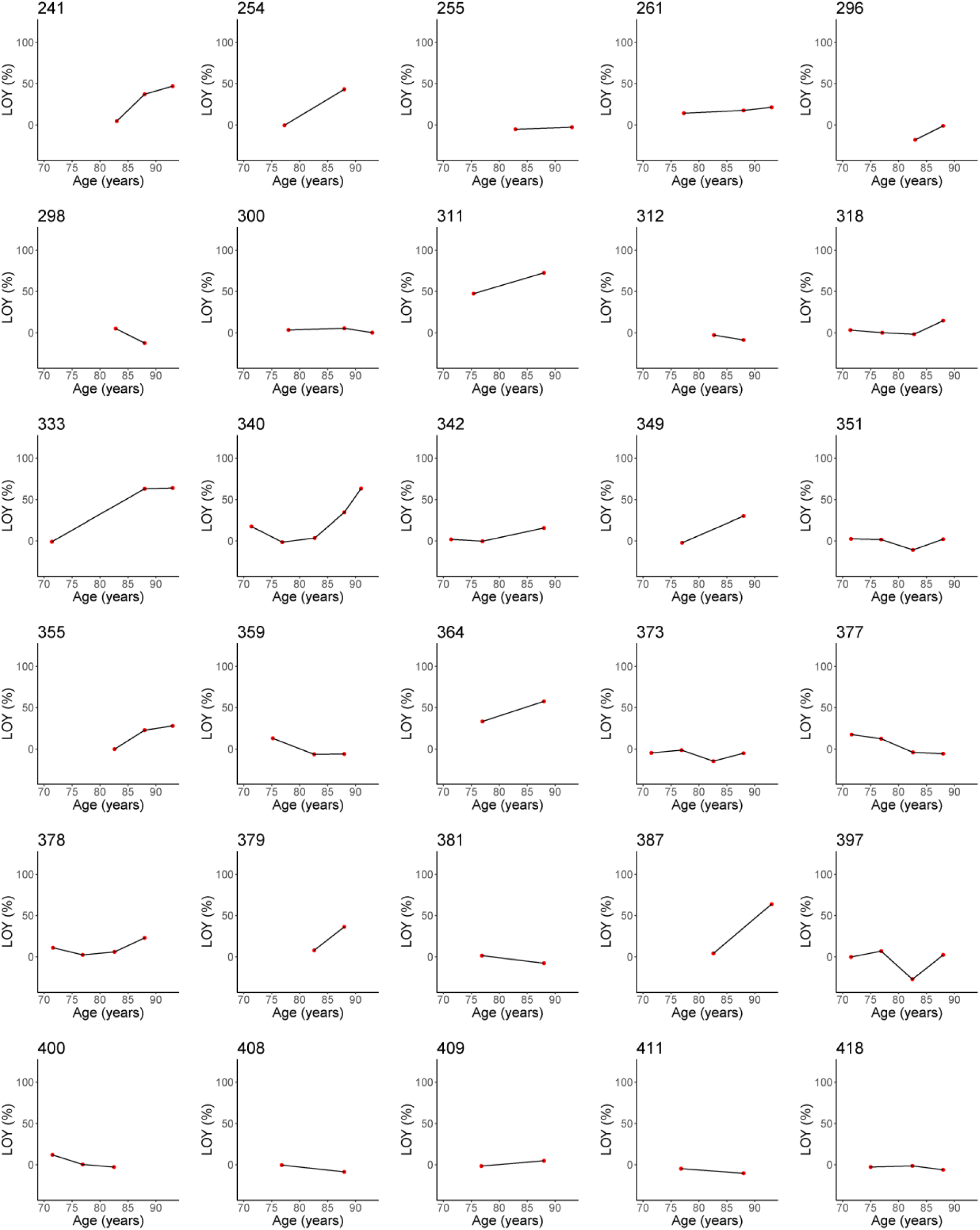
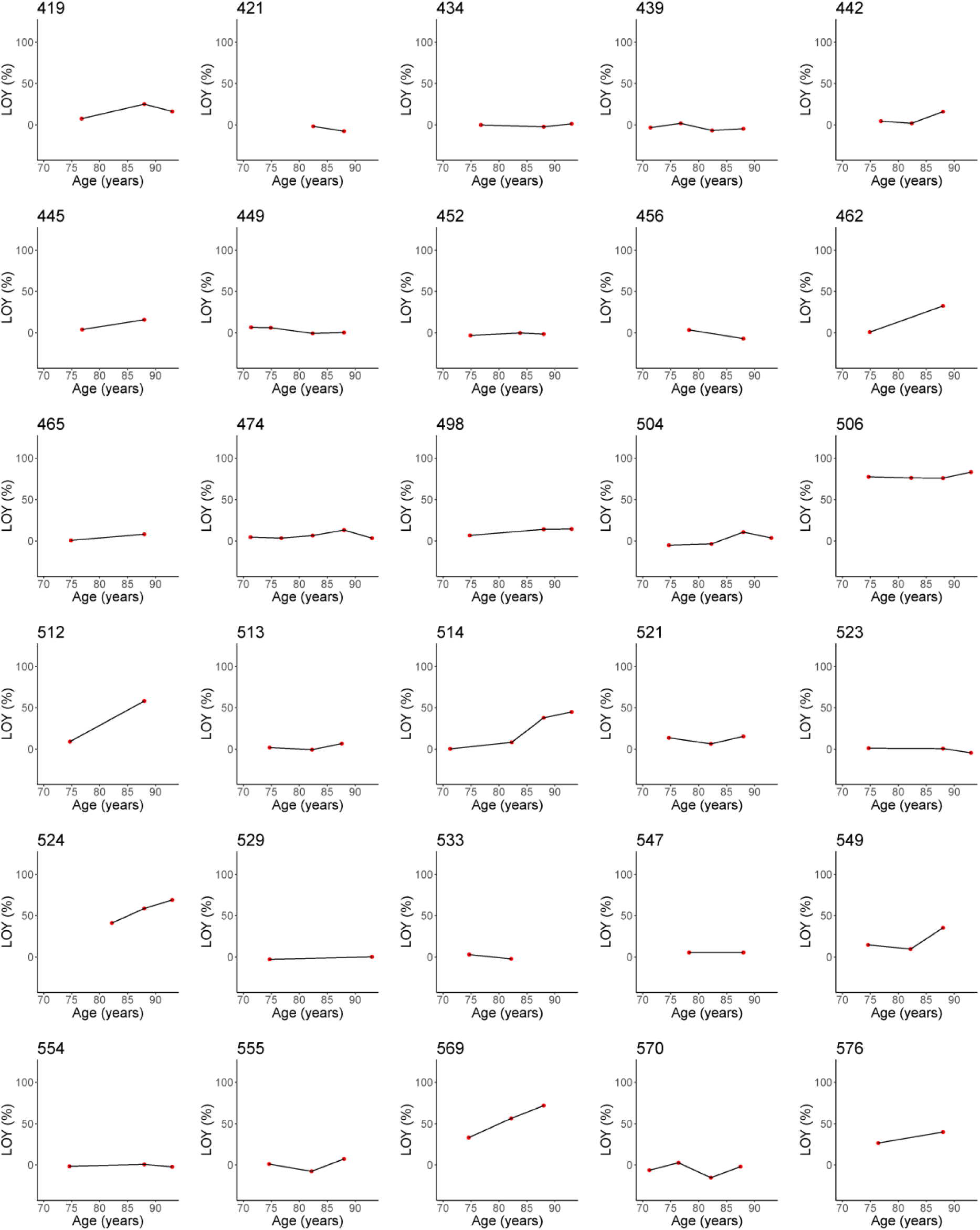
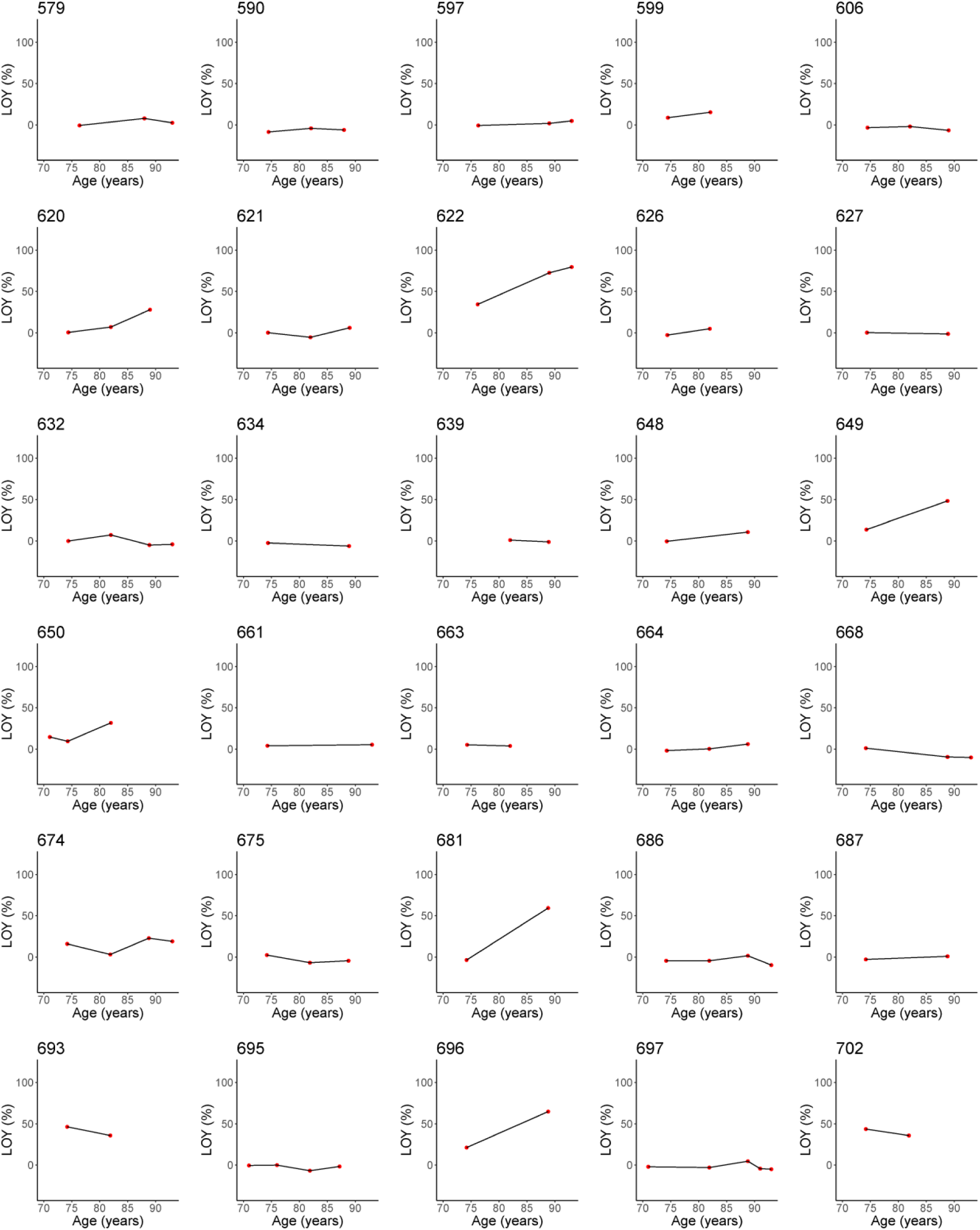
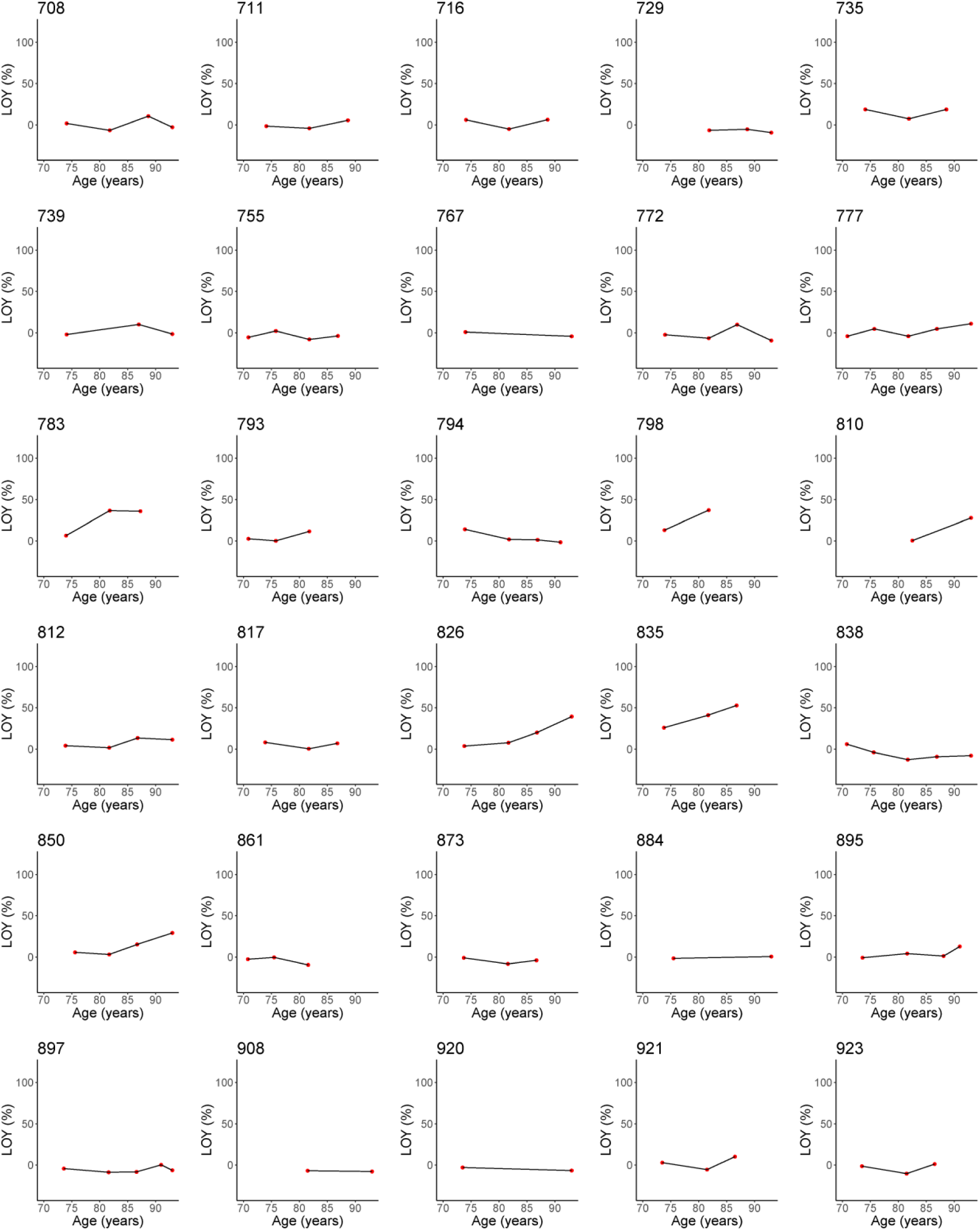
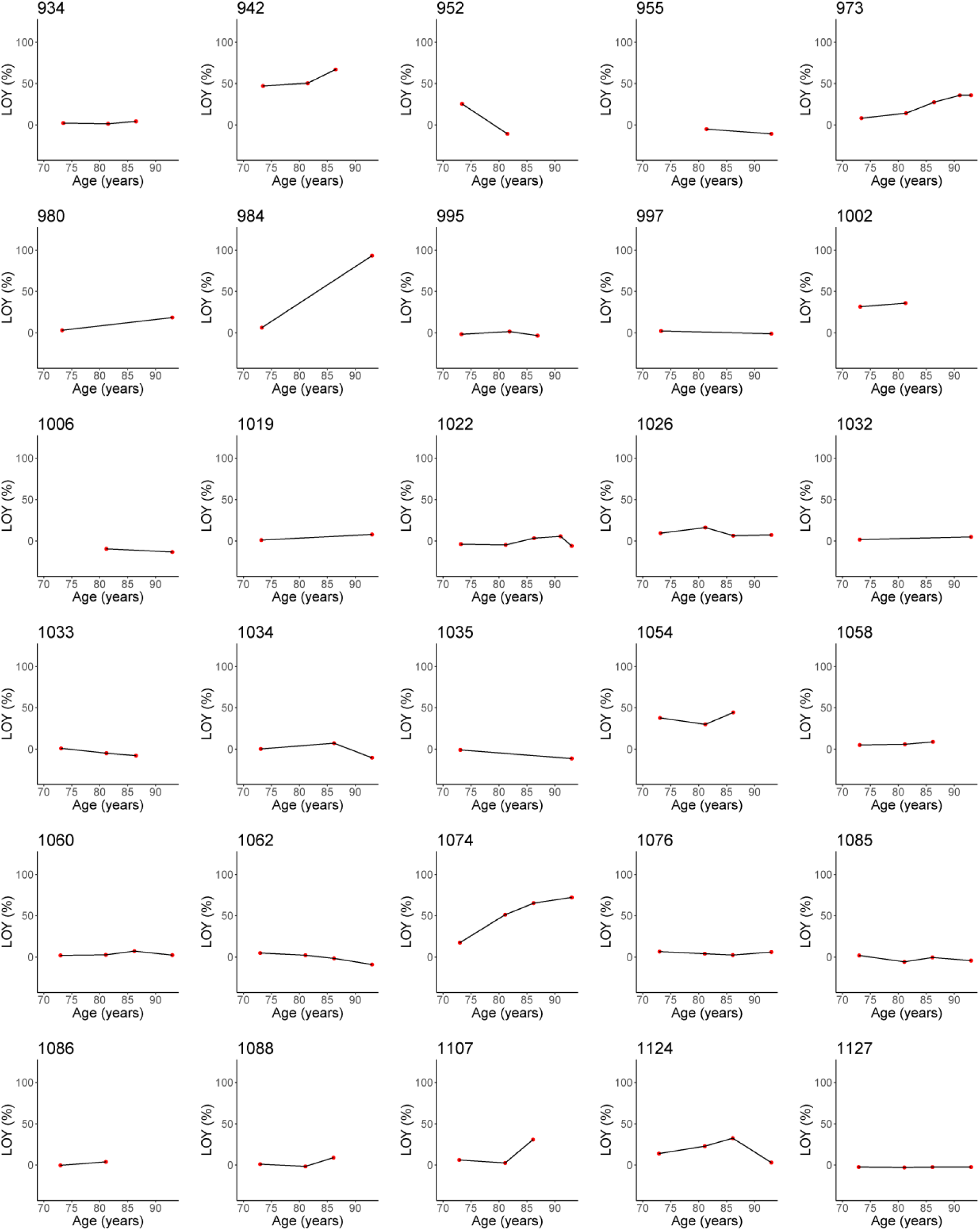
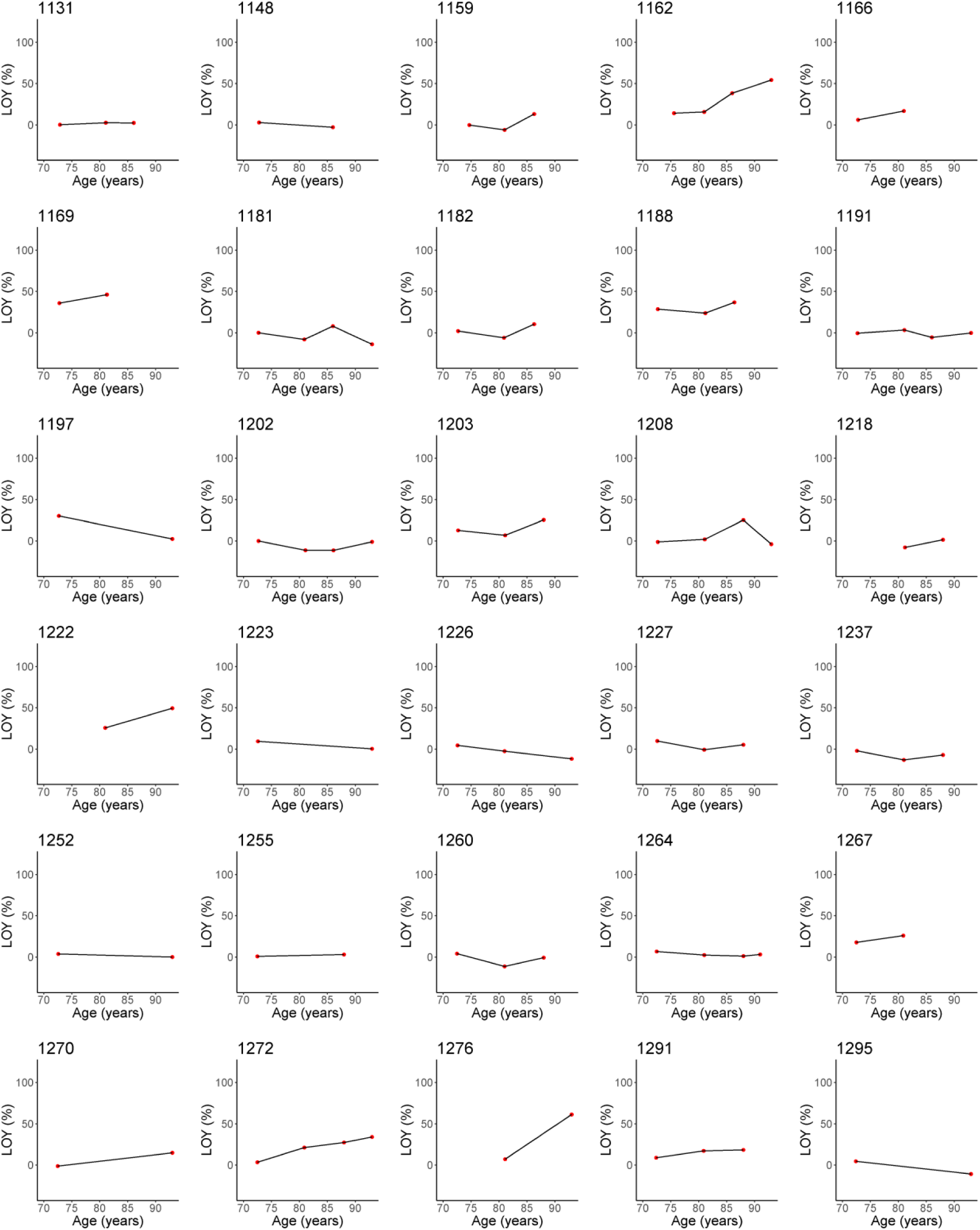
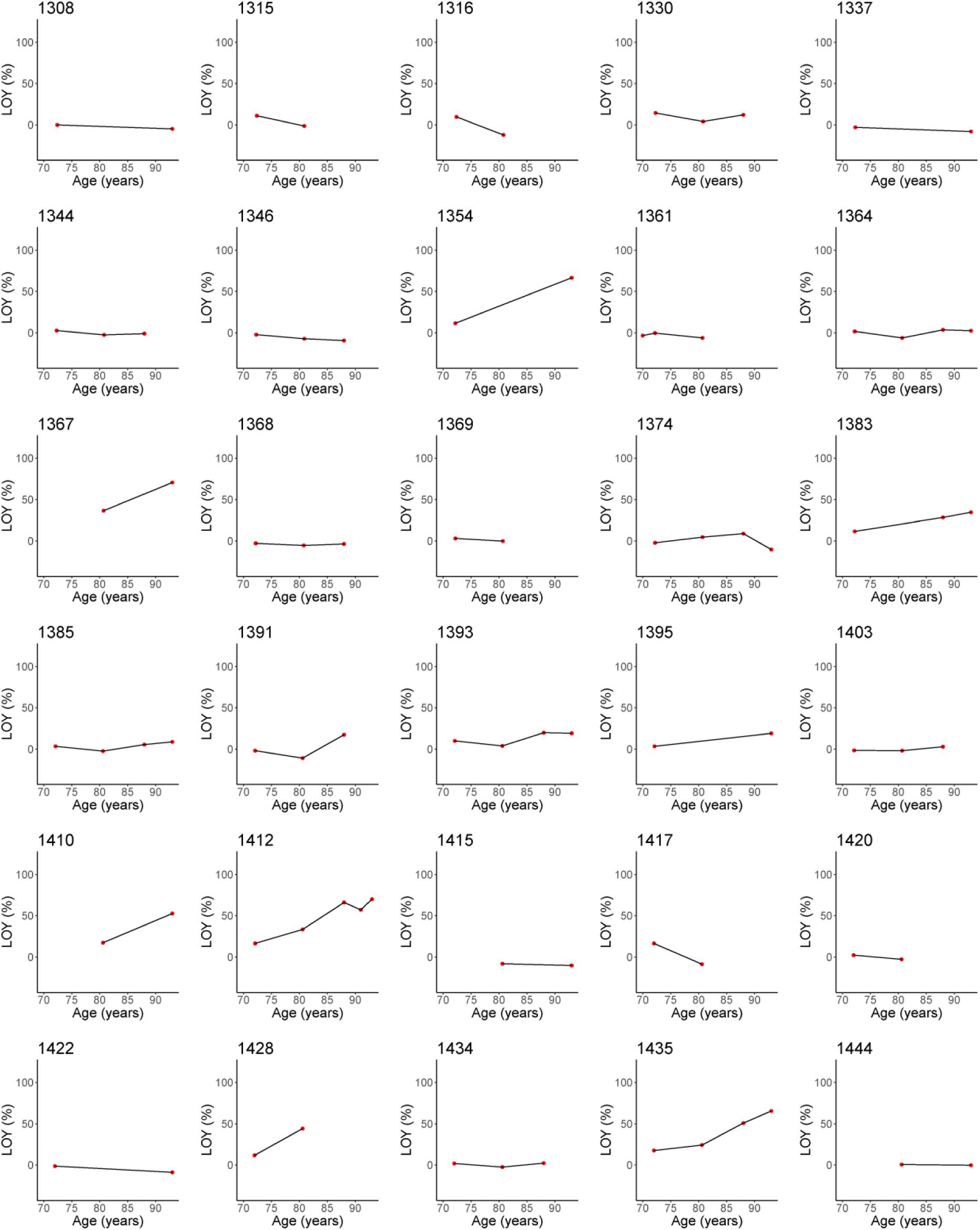
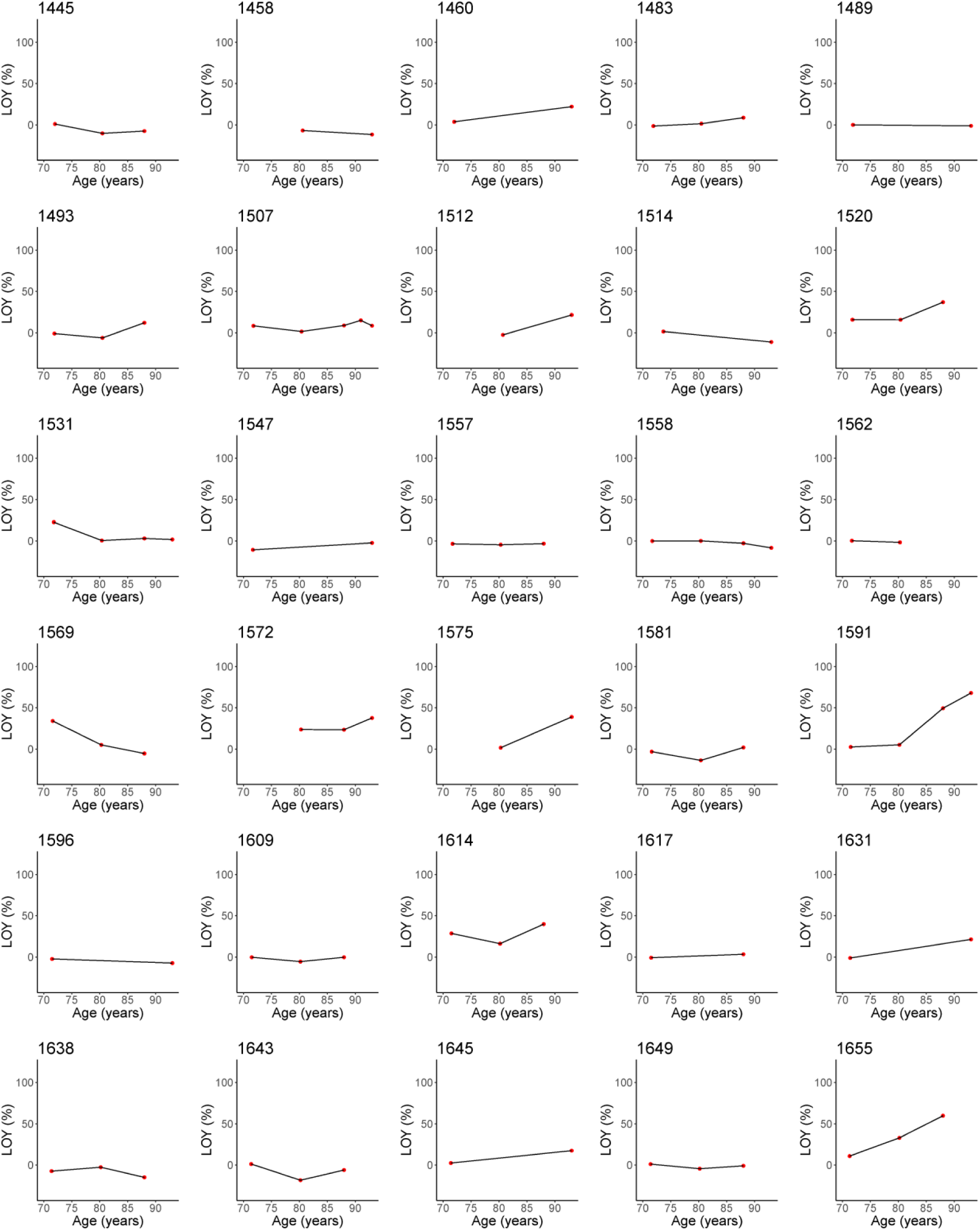
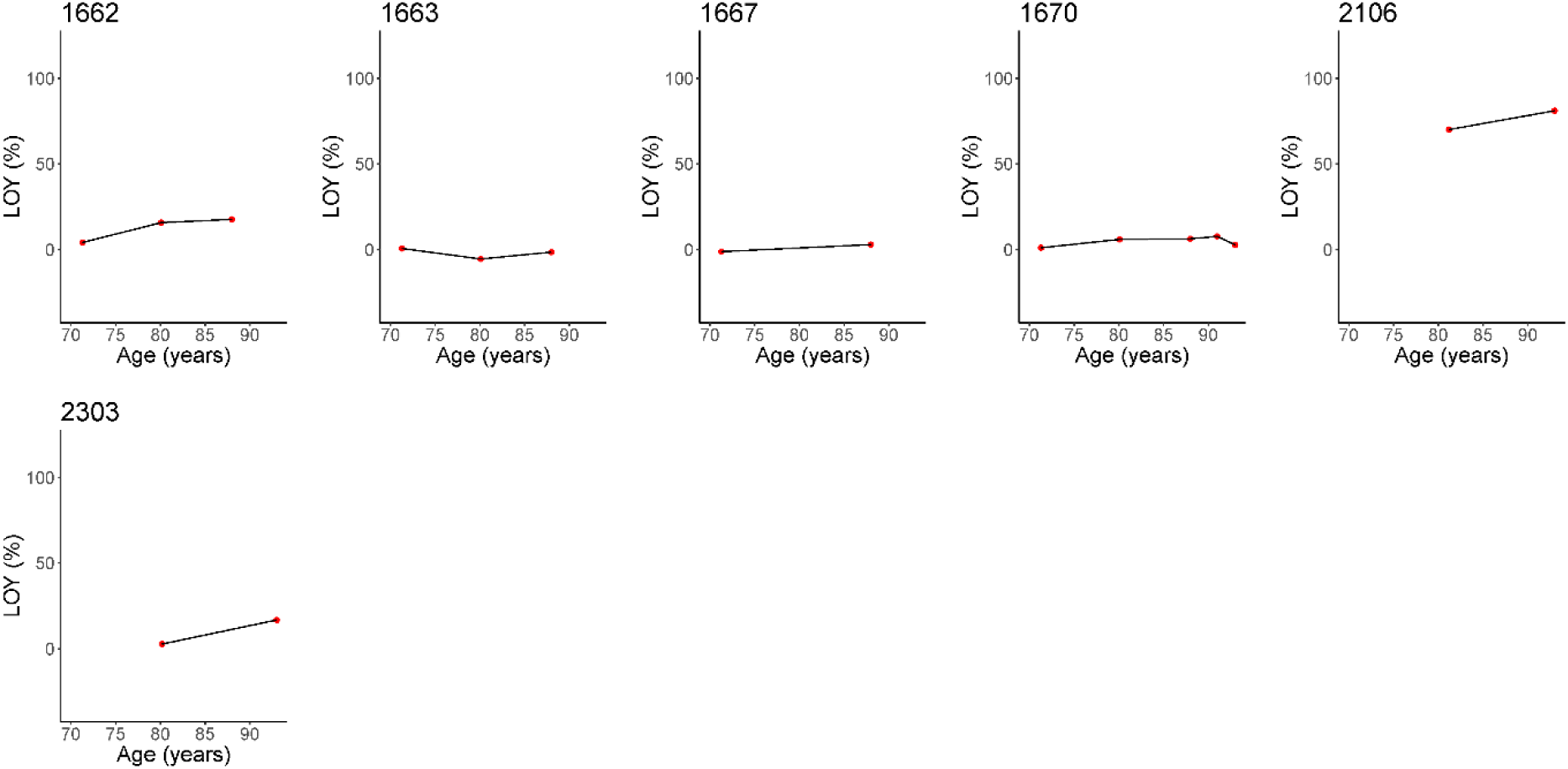
Intra-individual LOY dynamics in 276 individuals. Intra-individual LOY dynamics measured in 276 men visualized in LOY% from SNP-array genotyping data using a range of two to five LOY measurements for every individual. Plot numbering is according to the ID of individuals in the ULSAM cohort.

**Supplementary Table 1.**

| Method | Reference |
| --- | --- |
| FISH | Ganster, C. et al. New data shed light on Y-loss-related pathogenesis in myelodysplastic syndromes. *Genes Chromosomes Cancer* 54, 717-724 (2015). |
| FISH | Persani, L. et al. Increased loss of the Y chromosome in peripheral blood cells in male patients with autoimmune thyroiditis. *Journal of autoimmunity* 38, 1193-1196 (2012). |
| FISH | Lleo, A. et al. Y chromosome loss in male patients with primary biliary cirrhosis. *Journal of autoimmunity* 41, 87-91 (2013). |
| Karyotyping | Jacobs, P.A., Brunton, M., Court Brown, W.M., Doll, R. & Goldstein, H. Change of human chromosome count distribution with age: evidence for a sex differences. *Nature* 197, 1080-1081 (1963). |
| Karyotyping | UKCCG Loss of the Y chromosome from normal and neoplastic bone marrows. United Kingdom Cancer Cytogenetics Group (UKCCG). *Genes Chromosome Cancer* 5, 83-88 (1992). |
| qPCR | Noveski, P. et al. Loss of Y Chromosome in Peripheral Blood of Colorectal and Prostate Cancer Patients. *PLoS One* 11, e0146264 (2016). |
| qPCR | Kimura, A. et al. Loss of chromosome Y in blood, but not in brain, of suicide completers. *PLoS One* 13, e0190667 (2018). |
| qPCR | Hirata, T. et al. Investigation of chromosome Y loss in men with schizophrenia. *Neuropsychiatr Dis Treat* 14, 2115-2122 (2018). |
| SNP-array | Forsberg, L.A. et al. Mosaic loss of chromosome Y in peripheral blood is associated with shorter survival and higher risk of cancer. *Nat Genet* 46, 624-628 (2014). |
| SNP-array | Loftfield, E. et al. Predictors of mosaic chromosome Y loss and associations with mortality in the UK Biobank. *Sci Rep* 8, 12316 (2018). |
| SNP-array | Grassmann, F. et al. Y chromosome mosaicism is associated with age-1 related macular degeneration. *Eur J Hum Genet* this issue (2018). |
| SNP-array | Forsberg, L.A. et al. Mosaic loss of chromosome Y (LOY) in leukocytes matters. *Nature Genetics* In press (2018). |
| SNP-array | Dumanski, J.P. et al. Smoking is associated with mosaic loss of chromosome Y. *Science* 347, 81-83 (2015). |
| SNP-array | Wong, J.Y.Y. et al. Outdoor air pollution and mosaic loss of chromosome Y in older men from the Cardiovascular Health Study. *Environ Int* 116, 239-247 (2018). |
| SNP-array & qPCR | Haitjema, S. et al. Loss of Y Chromosome in Blood Is Associated with Major Cardiovascular Events during Follow-up in Men after Carotid Endarterectomy. *Circ Cardiovasc Genet* 10:e001544 (2017). |
| SNP-array & WGS | Dumanski, J.P. et al. Mosaic Loss of Chromosome Y in Blood Is Associated with Alzheimer Disease. *Am J Hum Genet* 98, 1208-1219 (2016). |
| SNP-array & WGS | Wright, D.J. et al. Genetic variants associated with mosaic Y chromosome loss highlight cell cycle genes and overlap with cancer susceptibility. *Nat Genet* 49, 674-679 (2017). |
| WGS | Zink, F. et al. Clonal hematopoiesis, with and without candidate driver mutations, is common in the elderly. *Blood* 130, 742-752 (2017). |

**Supplementary Table 2.**

| ID | LOY<br>estimated<br>by SNP- | LOY<br>estimated<br>by SNP- | LOY<br>estimated<br>by WGS | LOY<br>estimated<br>by ddPCR | ID | LOY<br>estimated<br>by SNP- | LOY<br>estimated<br>by SNP- | LOY<br>estimated<br>by WGS | LOY<br>estimated<br>by ddPCR |
| --- | --- | --- | --- | --- | --- | --- | --- | --- | --- |
| 63 | 0,01 | -1,64 |  | 4,75 | 1035 | 0,03 | -3,86 |  | 5,80 |
| 80 | -0,02 | 2,62 |  | 14,50 | 1060 | -0,07 | 8,96 |  | 18,85 |
| 91 | -0,08 | 11,03 |  | 15,40 | 1062 | 0,01 | -1,60 |  | 8,55 |
| 123 | -0,34 | 37,15 | 41,23 | 44,20 | 1074 | -0,97 | 74,09 | 67,35 | 67,35 |
| 153 | 0,03 | -3,75 |  | 5,65 | 1076 | -0,10 | 12,47 |  | 14,70 |
| 207 | 0,03 | -4,35 |  | -13,00 | 1085 | -0,02 | 2,76 |  | 9,30 |
| 241 | -0,51 | 50,40 | 53,34 | 55,35 | 1099 | 0,04 | -5,19 |  | 3,15 |
| 255 | -0,03 | 4,16 |  | 3,40 | 1103 | -0,19 | 23,46 |  | 36,05 |
| 261 | -0,22 | 26,66 | 30,86 | 32,85 | 1124 | -0,07 | 9,61 | 19,37 | 21,30 |
| 278 | -0,04 | 5,15 |  | 16,65 | 1127 | -0,03 | 4,50 |  | 5,00 |
| 300 | -0,05 | 7,02 |  | 12,35 | 1149 | 0,00 | -0,68 |  | 1,85 |
| 333 | -0,78 | 66,26 | 65,61 | 66,20 | 1162 | -0,61 | 57,30 | 59,84 | 64,00 |
| 355 | -0,29 | 32,77 | 35,49 | 38,65 | 1181 | 0,04 | -6,14 |  | 0,00 |
| 387 | -0,79 | 66,33 |  | 66,60 | 1191 | -0,05 | 6,63 |  | 0,00 |
| 419 | -0,18 | 21,90 | 27,93 | 31,10 | 1202 | -0,04 | 5,89 |  | 2,00 |
| 434 | -0,06 | 7,91 |  | 6,25 | 1208 | -0,02 | 3,26 | 8,64 | 13,15 |
| 474 | -0,08 | 10,04 | 4,39 | 6,00 | 1222 | -0,54 | 52,83 |  | 60,00 |
| 498 | -0,16 | 20,31 | 26,54 | 28,55 | 1223 | -0,05 | 7,05 |  | 17,30 |
| 504 | -0,08 | 10,30 | -0,47 | 1,65 | 1226 | 0,03 | -4,20 |  | 3,90 |
| 506 | -1,34 | 84,32 |  | 78,35 | 1252 | -0,05 | 6,82 |  | 18,45 |
| 514 | -0,48 | 48,60 |  | 54,75 | 1270 | -0,17 | 20,60 |  | 32,50 |
| 523 | -0,02 | 2,50 |  | 6,40 | 1272 | -0,35 | 38,53 | 42,73 | 43,35 |
| 524 | -0,90 | 71,20 | 67,92 | 71,45 | 1276 | -0,73 | 63,82 |  | 65,90 |
| 529 | -0,05 | 7,01 |  | 6,35 | 1295 | 0,02 | -3,27 |  | 4,20 |
| 554 | -0,03 | 4,71 |  | 10,35 | 1308 | -0,02 | 2,23 |  | 7,10 |
| 579 | -0,07 | 9,05 | 15,27 | 17,90 | 1337 | 0,01 | -0,81 |  | 3,60 |
| 597 | -0,09 | 11,20 |  | 21,45 | 1354 | -0,84 | 68,92 |  | 69,80 |
| 622 | -1,20 | 80,99 | 76,71 | 79,10 | 1364 | -0,07 | 9,13 |  | 22,80 |
| 632 | -0,02 | 3,11 |  | 8,35 | 1367 | -0,93 | 72,58 |  | 73,35 |
| 661 | -0,09 | 11,71 |  | 16,50 | 1374 | 0,02 | -2,71 |  | 7,55 |
| 668 | 0,02 | -2,67 |  | 7,55 | 1383 | -0,36 | 39,04 | 44,35 | 47,10 |
| 674 | -0,20 | 24,50 | 28,90 | 29,85 | 1385 | -0,12 | 14,93 |  | 22,55 |
| 686 | 0,02 | -2,22 |  | 1,20 | 1393 | -0,20 | 24,58 |  | 31,50 |
| 697 | -0,02 | 2,10 |  | 1,80 | 1395 | -0,20 | 24,52 |  | 37,80 |
| 708 | -0,03 | 4,05 | 3,72 | 5,25 | 1410 | -0,59 | 55,95 |  | 57,40 |
| 729 | 0,01 | -2,02 |  | 5,90 | 1412 | -0,92 | 72,15 | 72,40 | 74,25 |
| 739 | -0,04 | 5,43 |  | 3,40 | 1415 | 0,02 | -2,63 |  | 2,10 |
| 767 | -0,02 | 2,88 |  | 7,25 | 1422 | 0,01 | -1,43 |  | 1,15 |
| 772 | 0,01 | -1,91 |  | 6,95 | 1435 | -0,82 | 67,82 | 67,58 | 70,20 |
| 777 | -0,13 | 17,05 |  | 21,85 | 1444 | -0,05 | 6,62 |  | 15,45 |
| 810 | -0,29 | 32,69 |  | 37,90 | 1458 | 0,03 | -4,18 |  | -0,25 |
| 812 | -0,14 | 17,43 |  | 19,10 | 1460 | -0,23 | 27,34 |  | 38,15 |
| 826 | -0,41 | 43,46 | 45,97 | 47,00 | 1489 | -0,04 | 5,66 |  | 11,40 |
| 838 | 0,00 | -0,60 |  | 6,15 | 1507 | -0,12 | 14,78 |  | 22,70 |
| 850 | -0,30 | 34,03 |  | 39,80 | 1512 | -0,23 | 26,98 |  | 38,20 |
| 884 | -0,06 | 7,36 |  | 18,90 | 1514 | 0,03 | -3,58 |  | 6,50 |
| 896 | -0,19 | 22,97 |  | 32,05 | 1531 | -0,06 | 8,50 |  | 8,35 |
| 897 | -0,01 | 0,85 |  | 3,65 | 1547 | -0,03 | 4,72 |  | 9,35 |
| 908 | 0,00 | -0,50 |  | 4,50 | 1558 | 0,01 | -0,84 |  | 2,95 |
| 920 | 0,00 | 0,57 |  | 7,50 | 1572 | -0,39 | 41,95 | 45,21 | 46,35 |
| 955 | 0,02 | -3,39 |  | -0,50 | 1575 | -0,41 | 43,10 |  | 50,30 |
| 973 | -0,37 | 40,18 | 42,51 | 44,80 | 1591 | -0,87 | 70,08 | 68,30 | 69,35 |
| 980 | -0,20 | 24,05 |  | 22,60 | 1596 | 0,00 | 0,03 |  | 6,75 |
| 984 | -2,00 | 93,77 |  | 90,35 | 1628 | -0,01 | 0,78 |  | 6,20 |
| 997 | -0,04 | 5,79 |  | 4,00 | 1631 | -0,22 | 26,68 |  | 36,50 |
| 1006 | 0,04 | -5,51 |  | 3,00 | 1645 | -0,19 | 23,03 |  | 34,35 |
| 1019 | -0,11 | 14,26 |  | 7,10 | 1670 | -0,07 | 9,32 |  | 17,70 |
| 1022 | -0,01 | 1,38 |  | 10,50 | 1834 | -0,02 | 3,13 |  | 4,05 |
| 1026 | -0,11 | 13,74 |  | 27,05 | 2106 | -1,25 | 82,31 |  | 80,75 |
| 1032 | -0,09 | 11,48 |  | 21,20 | 2303 | -0,18 | 22,34 |  | 36,05 |
| 1034 | 0,02 | -3,01 |  | 3,20 |  |  |  |  |  |

**Supplementary Table 3.**

| Longitudinal Change | Individuals | frequency |
| --- | --- | --- |
| Unaffected | 185 | 67% |
| Increasing LOY (>1%LOY/year)* | 73 | 26% |
| Decreasing LOY (<1%LOY/year)* | 18 | 7% |

\*determined by the slope on a linear trendline

**Supplementary Table 4.**

| SNP-array platform | MSY<br>probes | PAR<br>probes |
| --- | --- | --- |
| 1MDuo | 4224 | 832 |
| 2.5M Omni | 2494 | 550 |
| Infinium QC Array 24 (XY chip) | 1401 | 545 |
| Omni Express Exome | 1387 | 809 |

**Supplementary Table 5.**

|  | Dilution<br>series 1<br>(N=4) | Dilution<br>series 2<br>(N=12) | Combined<br>series 1 and 2<br>(N=4+12=16) |
| --- | --- | --- | --- |
| Standard deviation | 0,027164 | 0,039043 | 0,036507 |
| Slope (b)* | 0,9466 | 0,9177 | 0,925 |
| LOD (ratio of cells with the Y chromosome) | 0,086089 | 0,127633 | 0,118401 |
| LOD (percent of cells with the Y chromosome) | 91,4 | 87,2 | 88,2 |
| LOD (LOY%) | 8,6 | 12,8 | 11,8 |

\*estimated by linear regression (supplementary figure 7)

